# Extracellular vesicles of a phytobeneficial bacterium trigger distinct systemic response in plant

**DOI:** 10.1101/2025.01.08.631853

**Authors:** Timothée Zannis-Peyrot, Lucas Degusseau, Pierre-Yves Dugas, Fabiola Bastian, Matthieu Gaucher, Vincent Gaillard, Gilles Comte, Florence Wisniewski-Dye, Isabelle Kerzaon, Céline Lavire, Ludovic Vial

## Abstract

Bacterial extracellular vesicles (EVs) are lipidic shuttles that play roles in virulence, inter-species competition, and in the induction of the host immune response. While they have primarily been investigated in animal-bacteria interactions, knowledge regarding phytobacterial EVs remains limited. Recent findings revealed that various biotic factors like hydroxycinnamic acids can regulate EVs production. Hydroxycinnamic acids, such as ferulic acid, are lignin components abundantly released in the plant environment, where they impact the ecology of numerous phytobacteria. *Azospirillum* sp. B510, a phytobeneficial bacterium, induces the accumulation of hydroxycinnamic acid derivatives in the plant and can metabolize them as carbon sources. We hypothesized that the presence of ferulic acid in the environment of *Azospirillum* sp. B510 would influence its EVs production in terms of size, quantity, and cargo. Conversely, we also proposed that EVs from this phytobacterium would influence plant metabolites and defense gene expression. Our results show both that ferulic acid (mimicking the plant environment) influences the content of EVs released by *Azospirillum* sp. B510 and that bacterial EVs also impact plant physiology at a systemic level according to their cargoes. This research provides the first evidence of a global effect of bacterial EVs on the plant and highlights the dynamics of plant-bacteria interactions mediated by EVs.

## Introduction

The plant environment is colonized by widely diverse microorganisms, and a dynamic molecular dialogue between the different partners enables the establishment of relationships, beneficial or not, throughout the plant’s life. On the plant side, the roots produce numerous primary and specialized (secondary) metabolites in the rhizosphere (Massalha *et al*., 2017). These metabolites, with their diverse functions, serve as pivotal factors that can influence the rhizosphere microbiome by modulating its growth environment. While some plant specialized metabolites can act as chemical defense molecules against pathogens, some others favor the development of beneficial bacteria (Nakayasu *et al*., 2023). The production of specialized metabolites is influenced by the rhizosphere microbiome, leading to a constant modulation of their accumulation. On the other hand, bacteria emit numerous molecular signals directly towards the plant facilitating colonization of the rhizosphere and thus the establishment of a sustainable ecological niche (Green and Mecsas, 2016). The interaction between microorganisms and plants is a highly intricated process, occurring via a sophisticated communication network and bacteria employs a multitude of mechanisms to colonize and survive in the plant environments (Trivedi *et al*., 2020). Among them, it has been recently proposed that bacterial extracellular vesicles (EVs) could play a significant role in this communication with its host plant (McMillan and Kuehn, 2021).

While several bacterial extracellular vesicles types have been described thus far, the most exhaustively studied are outer-membrane vesicles (OMVs). These mono-layer lipidic capsules are formed by blebbing of the outer membrane of gram-negative bacteria and contain periplasmic and cytoplasmic material (Toyofuku *et al*., 2019). Bacterial EVs cargoes are extremely diverse, and their biological significance is becoming increasingly complex. Indeed, they are involved in the transport of toxins, virulence factors, signal molecules, nucleic acids (DNA/RNA) (Toyofuku *et al*., 2023). As such, EVs derived from animal pathogens have been shown to play different roles in bacterial host colonization and host immunity modulation (Rosen *et al*., 1995; Kolling and Matthews, 1999; Bauman and Kuehn, 2006; Koeppen *et al*., 2016).

Despite the growing number of studies describing the roles of bacterial EVs in relation between animals and bacteria, little is known about their implications in plant-bacteria interactions. The majority of research into EVs from plant-associated bacteria has focused on phytopathogenic bacteria. For instance, EVs from *Pectobacterium* spp. can cause macerations of plant tissues due to the presence in their cargo of enzymes involved in cell-wall degradation (Jonca *et al*., 2021; Maphosa *et al*., 2023). *Pseudomonas syringae* EVs were shown to be produced *in vivo* during infection of *Arabidopsis thaliana* and may contribute to the virulence of *P. syringae*, as they harbor virulence factors (Janda *et al*., 2023). Phytopathogenic bacteria EVs have also been shown to allow the transport of Microbe-associated Molecular Patterns (MAMP) like Ef-TU and flagellin that can be recognized by plant immune receptors and therefore modulate the plant immune system (Tran *et al*., 2022; Chalupowicz *et al*., 2023). Interestingly, such modulation seems to be dependent on the bacteria from which EVs originate. Indeed, EVs from *P. syringae* and *Pseudomonas fluorescens* have been demonstrated to induce long term immunities against phytopathogens in *A. thaliana* but involving distinct immunity pathways (McMillan *et al*., 2021). It has also been shown that, within a given bacterium, changes in biotic or abiotic factors modulate the number and cargo of produced EVs. For example, a plant apoplast-mimicking minimal medium induced the accumulation of virulence factors in *P. syringae* EVs (McMillan and Kuehn, 2022). Similarly, the presence of genistein to mimic symbiosis conditions alters protein and lipid cargo of EVs extracted from the symbiotic bacteria *Sinorhizobium fredii* (Li *et al*., 2022). The presence of lignin derivatives in the environment of *Pseudomonas putida* has been linked to the production of bigger EVs and their enrichment in proteins involved in the degradation of plant cell wall compounds like hydroxycinnamic acids (HCAs) (Salvachúa *et al*., 2020).

While the effect of the plant environment on the content of phytobacterial EVs is increasingly well described, few studies have come full circle by investigating the effect of these modulated vesicles on plant models. We hypothesized that EVs from phytobeneficial bacteria also have an impact on plant physiology and conversely the bacterial EV production and/or cargoes are greatly influenced by the plant. For this purpose, we used ferulic acid (FA), a well-described HCAs, to mimic the plant environment. HCAs and their derivatives such as hydroxycinnamic acid amides (HCAAs) are specialized metabolites involved in the plant defense but also in the biosynthesis of plant cell walls as lignin precursors (Zeiss *et al*., 2021). As such, they are largely released in the plant environment where they play a major role in plant-bacteria interactions (Bhattacharya *et al*., 2010). HCAs, notably FA, have an anti-bacterial role due to their toxicity but can also be chemoattractants and used as trophic resources by some bacteria living in the rhizosphere (Herald and Davidson, 1983; Overhage *et al*., 1999). Hence, we used the well described plant growth-promoting rhizobacteria *Azospirillum* as a bacterial model. The phytobeneficial *Azospirillum* sp. B510, triggers the accumulation of HCAAs by the plant that can then be used as a carbon source by this bacterium (Valette *et al*., 2020). Moreover, this bacterium has been shown to induce systemic resistance in various plant models as in *Solanum lycopersicum* where it promotes resistance against *P. syringae* and *Botrytis cinerea* (Fujita *et al*., 2017). In this study, we show that the presence of HCAs in the environment of *Azospirillum* sp. B510 influences its EVs content. Furthermore, these qualitatively different EVs induce different molecular responses in *S. lycopersicum* at a systemic level. This molecular response varies according to the time elapsed after EVs inoculation. These results provide exciting clues to the existence of an EVs-mediated dynamism in the relationship between phytobacteria and the plant host.

## Materials and methods

### 2.1 Bacterial strain and growth conditions

*Azospirillum* sp. B510 was cultivated in LBm rich medium (yeast extract 5 g.L^−1^, peptone 10 g.L^−1^, NaCl 5 g.L^−1^) or AB minimal medium (Cangelosi *et al*., 1991) supplemented with succinate (100 mM) as a carbon source under agitation (180 rpm) when liquid medium was used or onto agar plates (agar 15 g.L^−1^). AB minimal medium was supplemented with ferulic acid (FA) (500 μM) when needed.

### 2.2 Analysis of ferulic acid degradation by *Azospirillum* sp. B510

Ferulic acid degradation by *Azospirillum* sp. B510 was tested as described in Campillo et al., 2014. Cells from 1 L of culture were pelleted and the supernatant from 15h to 22h of growth time was analyzed by UHPLC-DAD with an Agilent 1200 series HPLC system (Agilent Technologies, CA, USA) coupled to a UV-visible light (UV-vis) diode array detector (DAD) (Agilent 1200 series) and equipped with a Kromasil 100-5 C_18_ column (5 µm particle size; 250 × 4.6 mm; AkzoNobel).

Analyses were done at a flow rate of 1 mL.min^−1^ with an acidified (0.4% [*v/v*] formic acid in each solvent) MeOH/H_2_O gradient starting at 30:70 for 3 min, going to 39:61 in 3 min, to 100:0 in 4 min and maintained for 3 min before returning to the starting conditions in 4 min and equilibration for 1 min. Methanolic solutions (500 µM) of commercial ferulic acid, vanillic acid, and protocatechuic acid (Sigma Aldrich, MI, USA) or homemade 4-hydroxy-3-methoxyphenyl-β-hydroxypropionic acid, 4-hydroxy-3-methoxyphenyl-β-ketopropionic acid were also analyzed as references for UV-vis spectra and retention times. The injection volumes were 10 µL for culture medium and 5 µL for standard solutions. All data acquisitions and data reworks were done at 280 nm using Agilent Chemstation software B.03.02 (Agilent Technologies, CA, USA). FA degradation in the growth medium was verified by UHPLC-DAD before each EVs extraction.

### 2.3 Isolation of *Azospirillum* sp. B510 EVs (including protein and lipid quantification)

Bacterial cells grown for 20h were pelleted from two liters of culture in a Beckman Avanti J-E series centrifuge (JLA-14 rotor; 12,000 x *g*; 10 min). The supernatant was collected and filtered (0.45 µm HV, Millipore Durapore) to remove any residual bacterial cells. EVs were then concentrated using tangential flow with a 100 kDa cutoff filter (Sartorius). EVs were pelleted from the concentrate in a Beckman Optima LE-80K ultracentrifuge (vTi 65.2 rotor; 200,000 x *g*; 2h30; 4°C). The EVs pellet was resuspended in 3.5 mL of sterile dH_2_O then layered on top of an OptipPrep gradient (50%, 40%, 30%, 20%; 300 µL each). Gradients were centrifuged in a Beckman Optima LE-80K ultracentrifuge (vTi 65.2 rotor; 150,000 x *g*; 18 h; 4°C). Fractions were collected from the top of the ultracentrifuge tube. Optiprep was removed by washing in 4.7 mL sterile dH_2_O in a Beckman Optima LE-80K ultracentrifuge (vTi 65.2 rotor; 200,000 x *g*; 4h; 4°C). The supernatant was discarded and pellets were resuspended in 2 mL sterile dH_2_O before protein and lipid quantification and 10 µL of each sample were plated onto LB agar medium to verify the absence of contamination. EVs samples were used immediately for plant experiments or stored at −80°C before molecular content analysis.

For BCA microplate assays, 20 µL of EVs samples were mixed with 5 µL of 5% sodium deoxycholate (*w/v*) for 5 min to disrupt EVs membranes and added to 200 µL of BCA Working Solution. Absorbance was read at 562 nm using a plate reader (TECAN, Männendorf, Switzerland) and compared to a standard curve to calculate total protein concentration. For FM4-64 microplate assay, 10 µL of EVs samples were mixed with 200 µL of FM4-64 (1 µg.mL^−1^) during 30 min at 28°C in the dark. Fluorescence was measured in a plate reader (515 nm excitation, 635 nm emission; TECAN, Männendorf, Switzerland) and compared to a standard liposome curve prepared as described in Schaack *et al*., 2024. Each sample was measured as triplicate.

### 2.4 Scanning Electron Microscopy

Two milliliters of cells isolated from cell cultures (used for EVs extraction) were centrifuged at 600 x *g* for 3 min. The supernatant was discarded and cells were resuspended in 50 µL of Phosphate Buffer (PB) 0.1 M pH 7.4. Samples were fixed during 3h in PB containing 2% glutaraldehyde (*v/v*) and then filtered through a 0.2 µm filter. Samples were washed 3 times for 20 min in PB 0.2 M prior to post-fixation in 1% of osmium tetroxide (*v/v*) (diluted in PB 0.1 M) for 45 min. Samples were rinsed several times with water then dried overnight in 20% acetone (*w/v*) in a vacuum bell. Samples were incubated in 100% acetone for one hour at room temperature before being dried with Critical Point Dryer Leica EM CPD300 (Leica Microsystems, Wetzlar, Deustchland) (=V/med/120s/5/12/med/slow 100%) for 2h. Samples were mounted on stubs and sputter-coated with carbon, and then analyzed in a scanning electron microscope model FEG Merlin COMPACT VP (Carl Zeiss, Oberkochen, Deutschland).

### 2.5 Transmission Electron Microscopy

10 µL of EVs-containing samples were dropped on a hydrophilized copper Formvar grid (400 mesh) for 2 min. Fixed EVs were then contrasted with 1% tungsten disilicide (*w/v*) for 1 min before the grills got washed during 1 min with H_2_O and left to dry. Samples were analyzed using the transmission electron microscope model JEM −1400 Flash (Akishima, Tokyo, Japan).

### 2.6 Cryo Transmission Electron Microscopy

EVs suspensions were observed by cryo-TEM in order to preserve their morphology. 3 µL of the suspension were deposited onto a 300 mesh Lacey carbon film grid (EMS) and quench-frozen in liquid ethane using a Thermo Fisher Vitrobot. Samples were transferred in the microscope (JEOL 1400F - LaB6) using a precooled cryo-transfer holder (Fischione 2550) and observed with a Gatan Rio16 camera at an accelerating voltage of 120 kV in low-dose conditions.

### 2.7 Nanoparticles Tracking Analysis

EVs samples (3 biological replicates per condition) were diluted 1,000 times in dH_2_O and 1 mL of each resulting dilution was injected into the NTA (ZetaView, Particle Metrix GmbH, Germany). The zetaView software (8.05.16 SP3 version, ZetaView, Particle Metrix GmbH, Germany) was used to acquire and process each sample video. The instrument was set at temperature 25°C, sensitivity 80, 488 nm laser wavelength. Samples were analyzed in triplicates.

### 2.8 EVs protein sequencing and identification

EVs samples (3 biological replicates per condition) were processed following SP3 protocol (Hughes *et al*., 2019). Protein samples were reduced (TCEP, 5 mM final concentration, 45 min at 57°C) and alkylated (IAA, 10 mM final concentration, 45 min at ambient temperature) before being fixed on SP3 beads (10:1 bead/protein, incubation 5’ at 24°C, 1 000 rpm). After being washed thrice with EtOH 80% (*w/v*), beads-fixed proteins were digested in 96 µL of TEAB 100 mM with LysC/Trypsin (enzyme/protein ratio: 1/100) for 18 h at 37°C. Digested samples were then centrifuged (20,000 x *g*, 1 min) and supernatants containing peptides were recovered. Peptide concentration was determined for each sample by quantitative fluorometric peptide assay (Thermo Scientific) in order to inject 500 ng of peptides in the nanoLC-MS/MS system. Digested samples were analyzed by label free quantitative approach (LFQ) by nanoLC-MS/MS on Q Exactive HF high resolution mass spectrometer coupled with nanoUHPLC (Thermo Scientific). The three biological replicates for each condition were injected and loaded on a C18 Acclaim PepMap100 trap-column 300 µm ID x 5 mm, 5 µm, 100Å, (Thermo Scientific) for 3.0 minutes at 20 µL.min^−1^ with 2% ACN, 0.05% TFA in H_2_O and then separated on a C_18_ Acclaim Pepmap100 nano-column, 50 cm x 75 µm i.d, 2 µm, 100 Å (Thermo Scientific) with a 100 minutes linear gradient from 3.2% to 20% buffer B (A: 0.1% formic acid in H_2_O, B: 0.1% formic acid in ACN), from 20% to 32% of B in 20 min and then from 32% to 90% of B in 2 min, hold for 10 min and returned to the initial conditions in 2 min for 13 min. The total duration was set to 150 minutes with a flow rate of 300 nL.min^−1^ and the oven temperature was kept constant at 40°C. Samples were analyzed with a TOP15 DDA HCD method: MS data were acquired in a data dependent strategy selecting the fragmentation events based on the 15 most abundant precursor ions in the survey scan (350-1650 Th). The resolution of the MS scan was 120,000 at *m/z* 200 Th and for MS/MS scan the resolution was set to 15,000 at *m/z* 200 Th. Parameters for acquiring HCD MS/MS spectra were as follows: normalized collision energy = 27 and an isolation window width of 1.4 *m/z*. The precursors with unknown charge state, charge state of 1 and 8 or greater than 8 were excluded. Peptides selected for MS/MS acquisition were then placed on an exclusion list for 20 s using the dynamic exclusion mode to limit duplicate spectra. Raw data were processed with the Proteome Discoverer 2.5 (Thermo Scientific) software using SEQUEST HT search engine against Uniprot *Azospirillum* sp. B510 database (uniprot March 2023, 6,093 entries). Precursor mass tolerance was set at 10 ppm and fragment mass tolerance was set at 0.02 Da, and up to 2 missed cleavages were allowed. Oxidation (M), acetylation (Protein N-terminus) were set as variable modification and carbamidomethylation (C) as fixed modification. False discovery rate (FDR) of peptide identifications was calculated by the Percolator algorithm method, and a cut-off FDR value of 1 % was used. Protein quantification was done by Label Free Quantification (LFQ) approach, and LFQ abundance values were obtained and normalized to the total peptide amount. Protein quantitation was performed with precursor ions quantifier node in Proteome Discoverer 2.5 software with protein quantitation based on pairwise ratios and hypothesis t-test. Proteins present in the 3 biological replicates of at least one condition were considered as differentially expressed between the two conditions when FC > 2 or FC < 0.5 and *pv* < 0.05. Graphs were made using Rstudio (4.3.3) and GraphPadPrism 9 (9.1.0). Protein cellular localization prediction was done using CELLO (Yu *et al*., 2006). The mass spectrometry proteomics data have been deposited to the Center for Computational Mass Spectrometry repository (University of California, San Diego) via the MassIVE tool with the dataset identifier MassIVE MSMSV000096701.

### 2.9 Extraction of EVs metabolites and untargeted analysis

EVs samples (3 biological replicates per condition) were dried by lyophilization overnight. For each experiment, EVs powder samples were subjected to two successive extractions by adding twice 1 mL of MeOH 100% (*v/v*), and then twice 1 mL of MeOH/H_2_O 50:50 (*v/v*) with 15 min sonication at each extraction step. After centrifugation (10 min, 19,745 × *g*), the 4 supernatants obtained were pooled and transferred into a 5 mL hemolyze tube. The solvent was evaporated to obtain the crude extracts which were then solubilized at 10 mg.mL^−1^ in MeOH/H_2_O 80:20 (*v/v*) and stored at −80°C until further analysis. A quality control sample (QC) was done by mixing 10 µL of every sample used in the experiment. The analysis of *Azospirillum* sp. B510 EVs metabolites was performed by UHPLC-ESI-MS-QTOF on an UHPLC Agilent 1290 coupled to a UV-vis Diode Array Detector (DAD) (Agilent 1290 Infinity series) and a high-resolution Q-TOF 6546 mass spectrometer (Agilent Technologies, Santa Clara, CA, USA). Liquid chromatography was carried out using a Poroshell 120 EC-C_18_ column (3×100 mm, particle size 2.7 µm, Agilent technologies), preceded by a Poroshell C_18_ guard column (3×5 mm, particle size 2.7 µm, Agilent technologies) maintained at 40°C with an injection volume of 4 µL per sample. The mobile phase was a mixture of acidified acetonitrile (CH_3_CN) and acidified H_2_O (0.1% formic acid for each) applied at a flow-rate of 0.8 mL.min^−1^, and with a gradient (acidified H_2_O: acidified CH_3_CN, [*v/v*]) starting at 99:1 for 0.5 min, increasing to 70:30 in 9.5 min, going to 60:40 in 1.5, ramping up to 0.100 in 11.5 min and maintained for 1 min before returning to the starting conditions and equilibrating for 3 min. The QC sample was initially analyzed and then injected after every 6 samples in the run sequence to monitor the repeatability of the analysis. The quadrupole time-of-flight mass spectrometer (QTOF-MS) equipped with an electrospray ionization source (Dual AJS ESI Agilent, Santa Clara, CA, USA) was used in positive ionization mode for MS and MS^2^ analyses under the following conditions: drying gas (N_2_) flow of 12 L.min^−1^ at 320 °C, nebulizer pressure of 40 psi, sheath gas flow rate of 11 L.min^−1^ at 350 °C, with the capillary, nozzle and fragmentor voltages set to 3300 V, 500 V, and 100 V, respectively. The acquisition mass range was *m*/*z* 80 to 2,100 with a scan rate of 5 spectra.s^−1^, and the MS^2^ experiment was done with the collision energy set at 10, 20, 30, 40 or 50V. Reference solutions containing standard compounds (HP-0921, *m*/*z* 922.0098 and purine, *m/z* 121.0509) were constantly infused as an accurate mass reference. The UHPLC-ESI-MS-QTOF device was managed by the Agilent Mass Hunter DataAcquisition version 11.0 software. Data was processed using both workflow4metabolomics (Giacomoni *et al*., 2015) and MzMine 3.9 (Schmid *et al*., 2023). Specialized metabolites annotation was carried out both manually based on acquired spectral data and semi-automatically using Mass Hunter Qualitative Analysis B.07.00 software (Agilent Technologies) and SIRIUS with ZODIAC, CSI:FingerID and CANOPUS tools (Dührkop *et al*., 2015, 2019, 2021; Ludwig *et al*., 2019). Statistical analyses and graphical representations were performed using RStudio 4.3.1 (RStudio, PBC) and the package ‘ggplot2’ 3.4.

### 2.10 Extraction and analysis of EVs lipids

After addition of appropriate internal standards to the EVs samples (3 biological replicates per condition), total lipids were extracted twice with chloroform (CHCl₃)/ethanol (EtOH) (2:1, *v/v*). Lipid classes were separated by solid phase extraction (SPE). Briefly, NH_2_ cartridges (6cc.500 mg^− 1^, Waters) were washed with 8 mL hexane. Samples were loaded onto the cartridges. Then, they were washed with 20 mL CHCl₃ followed by 5 mL diethyl ether/acetic acid (98:2, *v/v*). Phospholipids were eluted with 2.5 mL MeOH/CHCl₃ (6:1, *v/v*) and then with a mixture of 0.05 M sodium acetate in MeOH/CHCl₃ (6:1, *v/v*). The fraction of total lipids and phospholipids were transmethylated with boron trifluoride in MeOH (14%). Transmethylation was carried out at 100 °C for 90 min. The derivatized fatty acid methyl esters were then extracted twice with isooctane and analyzed by Gas Chromatography coupled to a Flame Ionization Detector (GC-FID) using an HP6890 instrument equipped with a fused silica capillary BPX70 SGE column (60 m x 0.25 mm). The vector gas was hydrogen. Temperatures of the flame ionization detector and the split/splitless injector were set at 250 °C and 230 °C, respectively. Detected fatty acids were compared to standard molecules. Phospholipids were also analyzed by LC-MS/MS using a WATERS ACQUITY UPLC BEH Amide (130Å, 1.7 µm, 2.1 mm X 100 mm) column and an CH_3_CN/H_2_O gradient containing 10 mM ammonium formate. The flow rate was 0.6 mL.min^−1^. The HPLC system was coupled to a triple quadrupole mass spectrometer (AB SCIEX QTRAP 4500) equipped with electrospray ionization source (ESI). Source temperature was set at 500 °C, curtain gas at 40, gas 1 at 20 and gas 2 at 50, using nitrogen. Spray voltage was at −4,500 V. Analyses were achieved in negative mode using multiple reaction monitoring.

### 2.11 Growth conditions of *S. lycopersicum* and root inoculation

Seeds of *Solanum lycopersicum* var. Marmande were washed twice in sterile dH_2_O for 5 min, surface disinfected for 10 min in 2.5% sodium hypochlorite (*w/v*) and rinsed 4 times for 5 min in sterile dH_2_O. After disinfection, each seed was deposited on the surface of plant agar (6 g.L^−1^) poured into an 1.5 mL bottom-cut eppendorf tube (1.3 mL per tube). Seeds were left to germinate in the dark at 24°C for 4 days. Bottom-cut eppendorfs containing germinated seed were then transferred at the top of 15 mL Falcon tubes containing 13 mL of Murashige and Skoog ½ strength pH 5.8 sterile medium and let to grow for 14 days in a growth chamber (photoperiod 16 h of day, temperature 24°C day and 22°C night, 60% to 70% humidity, medium refilled when needed). After 14 days of growth, roots were inoculated with either sterile water or 1.10^9^ EV particles (13 µg proteins) (Control, EV or EV FA condition; 120 plants per condition) and put back in the growth chamber (**Supp. Fig. 1**). An aliquot of the plant growth medium was plated onto LB agar plates to assess the sterility of the experimental conditions.

### 2.12 Extraction of *S. lycopersicum* metabolites

Eighteen hours and five days post-inoculation, 60 plants of each condition were randomly gathered in 6 groups to reduce variability prior to roots and aerial parts harvest. Roots and aerial parts were harvested separately before being quickly frozen in liquid nitrogen for further analyses. After harvest, plant parts were dried by lyophilization overnight then crushed with stainless steel grinding balls (FastPrep-24 5G, MP Biomedicals, USA). For each experiment, plant powder samples were subjected to two successive extractions by adding twice 1 mL of acidified MeOH/H_2_O 80:20 (*v/v*) (0.1% formic acid), and then twice 1 mL of acidified MeOH 100% (0.1% formic acid) with 15 min sonication at each extraction step. After centrifugation (10 min, 19,745 ×*g*), the 4 supernatants obtained were pooled and transferred into a 5 mL hemolyze tube. The solvent was evaporated to obtain the crude extracts which were then solubilized at 10 mg.mL^−1^ in MeOH/H_2_O 80:20 (*v/v*) and stored at −80°C until further analysis. For each experiment, a QC was done by mixing 6 µL of every sample used in the experiment.

### 2.13 Targeted analysis of free amino acid content of *S. lycopersicum*

Tomato free amino acids were analyzed as described previously (Henderson et *al.*, 2000). The analysis was performed on an HPLC Agilent 1100 coupled to a DAD and a fluorometric detector (FLD) (Agilent 1100 Infinity series). Norvaline (2 µL, 2.5 mM) was used as an internal standard and added to 100 µL of the tested samples. Briefly, liquid chromatography was carried out using a Poroshell 120 EC-C_18_ column (3×100 mm, particle size 2.7 µm, Agilent technologies), preceded by a Poroshell C_18_ guard column (3×5 mm, particle size 2.7 µm, Agilent technologies) maintained at 45°C with an injection volume of 3 µL per sample. A mix of 24 amino acids (α-aminobutyric acid, alanine, arginine, asparagine, aspartic acid, citruline, γ-aminobutyric acid, glutamine, glutamic acid, glycine, histidine, isoleucine, leucine, lysine, methionine, hydroxyproline, ornithine, phenylalanine, proline, serine, threonine, tryptophan, tyrosine, valine) prepared as described previously (Henderson et *al.*, 2000) was injected as standard. The HPLC-UV/DAD-FLD device was managed by Mass Hunter Workstation Acquisition B.07.00 software, and the data was converted and reworked with Mass Hunter Qualitative Analysis B.07.00 software (Agilent Technologies). Statistical analyses and graphical representations were performed using RStudio 4.3.1 (RStudio, PBC) and the packages ‘mixOmics’ 6.24 (Rohart *et al*., 2017), ‘ropls’ 1.32.0 (Thévenot *et al*., 2015) and ‘ggplot2’ 3.4.4.

### 2.14 Analysis of specialized metabolites content of *S. lycopersicum*

The analysis of tomato specialized metabolites was performed by UHPLC-ESI-MS-QTOF on an UHPLC Agilent 1290 coupled to a DAD (Agilent 1290 Infinity series) and a high-resolution Q-TOF 6546 mass spectrometer (Agilent Technologies, Santa Clara, CA, USA). Liquid chromatography was carried out using a Poroshell 120 EC-C_18_ column (3×150 mm, particle size 2.7 µm, Agilent technologies), preceded by a Poroshell C_18_ guard column (3×5 mm, particle size 2.7 µm, Agilent technologies) maintained at 45°C with an injection volume of 3 µL per sample. For tomato root metabolites, the mobile phase was a mixture of acidified CH_3_CN and acidified H_2_O (0.1% formic acid for each) applied at a flow-rate of 0.8 mL.min^−1^, and with a gradient (acidified H_2_O: acidified CH_3_CN, [*v/v*]) starting at 99:1 for 1.5 min, increasing to 90:10 in 1.5 min, going to 84:16 in 7 min, going to 82:18 in 2.5 min, going to 73:27 in 7.5 min and maintained for 2 min, then going to 68:32 in 1 min, going to 0:100 in 3 min and maintained for 2 min, before returning to the starting conditions and equilibrating for 2 min. For tomato aerial part metabolites, the same solvents were applied at a flow-rate of 0.8 mL.min^−1^ with a gradient (acidified H_2_O: acidified CH_3_CN, [*v/v*]) starting at 99:1 for 1.5 min, increasing to 93:7 in 1.5 min, going to 84:16 in 7 min, going to 82:18 in 2.5 min, going to 73:27 in 7.5 min and maintained for 2 min, then going to 68:32 in 1 min, going to 0:100 in 3 min and maintained for 2 min, before returning to the starting conditions and equilibrating for 2 min. For each experiment, the QC sample was initially analyzed and then injected after every 6 samples in the run sequence to monitor the repeatability of the analysis. The quadrupole time-of-flight mass spectrometer (QTOF-MS) equipped with an electrospray ionization source (Dual AJS ESI Agilent, Santa Clara, CA, USA) was used in positive ionization mode for MS and tandem MS analyses under the following conditions: drying gas (N_2_) flow of 12 L.min^−1^ at 320 °C, nebulizer pressure of 40 psi, sheath gas flow rate of 11 L.min^−1^ at 350 °C, with the capillary, nozzle and fragmentor voltages set to 3300 V, 500 V, and 150 V, respectively. The acquisition mass range was *m*/*z* 70 to 2,000 with a scan rate of 2 spectra.s^−1^, and the MS^2^ experiment was done on QC sample with the collision energy set at 10, 20, 30, 40 or 50V. Reference solutions containing standard compound (HP-0921, *m*/*z* 922.0098 and purine, *m/z* 121.0509) were constantly infused as an accurate mass reference. The UHPLC-ESI-MS-QTOF device was managed by Mass Hunter Workstation Acquisition B.07.00 software. Data was processed using both workflow4metabolomics for specialized metabolites profiles matrix obtention (Giacomoni *et al*., 2015). Specialized metabolites annotation was carried out semi-automatically based on spectral data acquired on MzMine 3.9 (Schmid *et al*., 2023) using SIRIUS with ZODIAC, CSI:FingerID and CANOPUS tools (Dührkop *et al*., 2015, 2019, 2021; Ludwig *et al*., 2019) and manually using Mass Hunter Qualitative Analysis B.07.00 software (Agilent Technologies). Statistical analyses and graphical representations were performed using RStudio 4.3.1 (RStudio, PBC) and the packages ‘mixOmics’ 6.24 (Rohart *et al*., 2017), ‘ropls’ 1.32.0 (Thévenot *et al*., 2015) and ‘ggplot2’ 3.4.4. The mass spectrometry metabolomicss data have been deposited to the Center for Computational Mass Spectrometry repository (University of California, San Diego) via the MassIVE tool with the dataset identifier MassIVE MSV000096676.

### 2.15 Analysis of defense gene expression in *S. lycopersicum* leaves

Each sample (biological replicate) corresponds to a bulk of the youngest expanded leaf from fifteen plants per treatment (Control, EV and EV FA). Hundred mg of leaf tissues were ground (Retsch) and total RNA was extracted with the Macherey-Nagel Nucleospin RNA Plant Kit (Macherey-Nagel) and by adding 2% PVP-40 in the extraction buffer. Reverse transcription was performed as described by Promega (M-MLV) and the complete absence of gDNA checked by polymerase chain reaction (PCR) using EF-α (elongation factor α) primers flanking an intron, as described in Gilliland et *al.*, 1990. Quantitative real-time PCR were performed using a patented set of primers for 28 defense genes and the 3 reference genes (Bernonville *et al*., 2011). Briefly 0.15 µl of the cDNA sample was mixed in a final volume of 15 µl with 7.5 µl of MESA BLUE qPCR MasterMix (Eurogentec), 1.5 µl of each primer (appropriately diluted) and 4.35 µl of water. Reaction was performed with a CFX96TM real time system (Bio-Rad) using the following program: 95°C for 5 min, 40 cycles comprising 95°C for 3 s and 60°C for 60 s, with real-time fluorescence monitoring. Relative changes in defense gene expression (log2 ratio) were calculated using the 2−ΔΔCT method (Schmittgen and Livak, 2008) and the 3 internal reference genes for normalization (Vandesompele *et al*., 2002). The value of CT-treated samples was used for the calculation of the log2 ratio of each defense gene. Data were obtained from the same 360 tomato plants used in the metabolomic experiment, grouped in four biological replicates per treatment (Control, EV and EV FA, 15 bulk leaves per condition) and per harvest time (18 h and 5 days post-inoculation). Statistical analyses and graphical representations were performed using RStudio 4.3.1 (RStudio, PBC) and the packages ‘mixOmics’ 6.24 (Rohart *et al*., 2017), ‘ropls’ 1.32.0 (Thévenot *et al*., 2015) and ‘ggplot2’ 3.4.4.

## Results

### 3.1 Degradation of ferulic acid and conditions for extraction of *Azospirillum* sp. B510 EVs

*Azospirillum* sp. B510 was known to use HCAAs as carbon sources (unpublished results) but its ability to degrade HCAs was not assessed until now. *Azospirillum* sp. B510 was able to grow in AB minimal media with ferulic acid (FA) or coumaric acid (CA) as the only carbon source (**Supp Fig. 2**). By comparison with other genomic regions involved in the degradation of FA and CA in bacteria related to *Azospirillum*, we identified a genomic region potentially involved in the degradation of HCAs : AZL_b03620 to AZL_b03690 (**Supp. Fig. 3**)

FA degradation in AB medium supplemented with succinate (ABS) was tested. In ABS supplemented with FA (500 µM), FA concentration dropped between 14h and 19h; after 20h of growth, FA was no longer detected. Vanillic acid (VA) was identified as one of the degradation intermediates of FA by *Azospirillum* sp. B510. It increased in the supernatant from 14h to 18h and decreased from 18h to 20h, probably feeding the protocatechuic acid (PCA) pathway (**Supp. Fig. 4**). Hence, after 20h of growth in ABS supplemented with FA, FA and VA were degraded by *Azospirillum* sp. B510, indicating that the enzymes responsible for the degradation of HCAs were produced and active. Thus, the incubation time chosen for all EVs extraction was set at 20h.

### 3.2 Visualization and quantification of *Azospirillum* sp. B510 EVs

Visualization by scanning electron microscopy (SEM) of *Azospirillum* sp. B510 cells grown for 20h in ABS minimal medium in the presence or absence of FA led to the observation of numerous spherical structures with some apparently blebbing from bacteria membranes and others seemingly free (**Fig. 1A**). In order to obtain EVs samples of sufficient quantity and quality, cultures cleared from bacteria were concentrated using tangential flow filtration and resulting concentrated EVs samples were further purified using gradient density ultracentrifugation. Each ultracentrifuge gradient layer was analyzed using Nanoparticle Tracking Analysis (NTA) and the protein and lipid content of these layers were also quantified. EVs were detected by NTA only in the 40% (ρ = 1.21 g.mL^−1^) Optiprep layer, confirming the indirect quantification results using BCA and FM4-64 assays (**Fig. 1E**, **Supp. Fig. 5**). The 40% Optiprep layers were subsequently observed using electron microscopy to confirm the presence of EVs and also to ensure the absence of contaminating material. No bacteria, pili, flagella or membrane debris were detected in transmission electron microscopy (TEM) and cryo-electron microscopy (Cryo-EM) samples, assessing the purity and quality of extracted EVs samples (**Fig. 1B**). The morphology of EVs after extraction was analyzed by Cryo-EM. As seen on image (**Fig. 1C**) that is representative of the observed samples by Cryo-EM in terms of EVs structure, the purification steps seemed to have no impact on the integrity of EVs membrane. As only one lipid membrane was observed in our micrographs, the large majority of purified *Azospirillum* sp. B510 EVs were OMVs. Lastly, to investigate the impact of FA on the production of EVs by *Azospirillum* sp. B510, EVs sizes were evaluated using NTA. Purified EVs were homogeneous with a mean size of 135 nm. The presence of FA in the growth medium of *Azospirillum* sp. B510 did not significantly affect the average size of produced EVs (**Fig. 1D-E**).

**Figure 1:**
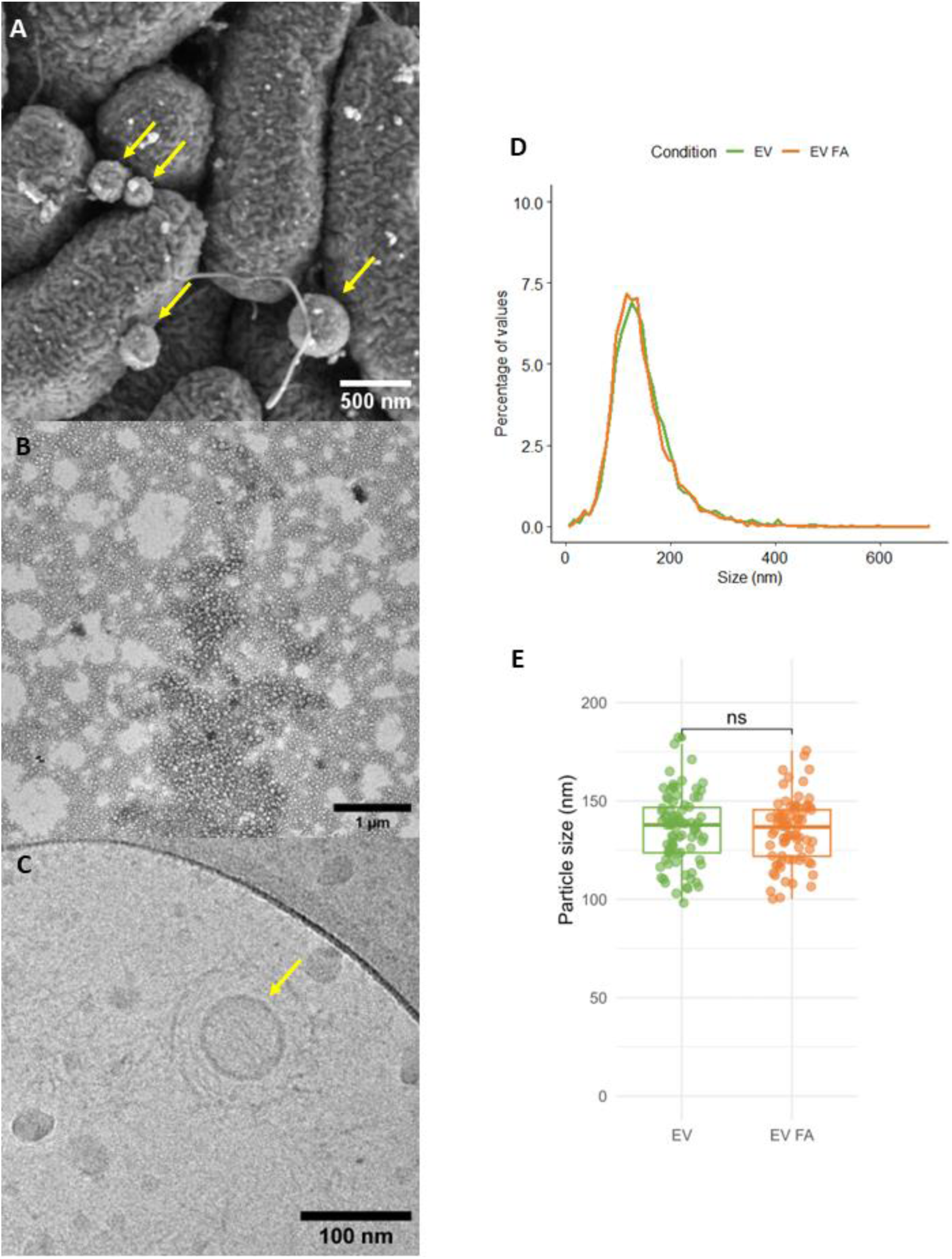
Visualization and size measurements of *Azospirillum* sp. B510 EVs. **(A)** SEM image of *Azospirillum* sp. B510 and its EVs grown in liquid AB minimal medium. **(B)** TEM image of purified EVs samples used as inoculum in plant experiments. **(C)** Cryo-EM of an EV from a purified EVs sample. For each microscopy technique, the depicted image is representative of at least 15 pictures. **(D)** Size distribution of *Azospirillum* sp. B510 EVs from FA free (EV) and FA containing (EV FA) culture obtained by NTA. **(E)** Mean size comparison of *Azospirillum* sp. B510 EVs from FA free (EV) and FA-containing (EV FA) culture obtained in GDUC 40% fraction by NTA (Student’s *t-test*, *pv* > 0.05, ns = not significant). NTA analyses were conducted on 3 biological replicates. Yellow arrow on SEM and Cryo-EM micrographs indicate EVs.

### 3.3 Effect of ferulic acid on the cargo composition of *Azospirillum* sp. B510 EVs

#### 3.3.1 Effect of ferulic acid on the proteinaceous cargo of *Azospirillum* sp. B510 EVs

The proteomic analysis of 3 biological replicates of *Azospirillum* sp. B510 EVs by a Q Exactive HF mass spectrometer allowed to identify 1,594 proteins in the EV condition out of the 6,309 proteins potentially encoded by the *Azospirillum* sp. B510 genome.

Amongst the 200 most abundant proteins of this EV cargo were found 52 ribosomal proteins and multiple known protein markers of bacterial EVs, such as the chaperonin GroEL, TolB, Ef-TU and components of the flagellar apparatus (**Supp. Table 1**). Adding FA to the growth medium induced the depletion of 60 proteins and the enrichment of 69 proteins in the *Azospirillum* sp. B510 EVs cargo. Only 2 proteins were solely found in the EV condition whereas 8 were unique to the EV FA condition. Among the depleted proteins, 4 flagellar apparatus proteins were found, of which AZL_e01020 (FliC). Moreover, multiple receptors and transporters involved in the transport of nutrients such as AZL_018060 (ProX), 2 putrescine binding proteins AZL_c00220 and AZL_c00690, and transporters of metal ions such as AZL_007910 and AZL_004770 were less abundant in the EV condition than in the EV FA condition. Lastly, 14 yet uncharacterized proteins were found depleted in the EV condition compared to the EV FA condition (**Fig. 2A**, **Table 1**). *Azospirillum* sp. B510 EV FA annotated proteins were depleted in the category “transport” of the Biological Processes GO, from which choline transport, organic acid and organic substance transport, and amino acid transport. They were also depleted in aspartate, carbohydrate and glucose metabolic processes, in cell adhesion and flagellum-dependent cell motility. On the opposite, *Azospirillum* sp. B510 EV FA were enriched in the GO categories 3,4-dihydroxybenzoate catabolic process, amino acid catabolic process, aromatic compound catabolic process and cellular aldehyde process. They were also enriched in proteins belonging to the GO categories of the cell cycle, fatty acid beta-oxydation, phosphorelay signal transduction system and P450-containing electron transport chain (**Fig. 2B**). Also, by using the classification of the KEGG pathway protein families, proteins of *Azospirillum* sp. B510 EV FA appeared enriched in families related to unclassified metabolism, carbohydrate metabolism, energy metabolism and amino acid metabolism and depleted in families corresponding to environmental information processing, genetic information processing, cellular processing and glycan biosynthesis and metabolism (**Fig. 2C**). The presence of FA in the growth medium induced the enrichment in *Azospirillum* sp. B510 EVs of Ech (AZL_b03680) and Vdh (AZL_b03690) involved in the degradation of FA to VA and PcaG (AZL_d00460), PcaH (AZL_d00470), PcaB (AZL_d00480) belonging to the *pca* operon (**Supp. Fig. 3**). Globally, FA induced the enrichment of proteins coded by genes mostly located on the replicon b of the composite genome of *Azospirillum* sp. B510 and especially from 2 gene clusters (**Fig. 2D**, **Supp. Fig. 6**).

**Table 1:** Differentially accumulated proteins in *Azospirillum* sp. B510 EVs in presence or absence of ferulic acid in growth medium. Proteins were found in 3 biological replicates of at least one growth condition.

| Log2FC<br>(EV FA / EV) | Accession | Gene ID | Description | Gene name | Unique peptides number | MW [kDa] |
| --- | --- | --- | --- | --- | --- | --- |
| -6,64 | D3P073 | AZL_a05210 | Beta-propeller repeat-containing protein |  | 5 | 34,9 |
| -6,64 | D3NVL5 | AZL_018060 | Glycine betaine/proline transport system substrate-binding protein | *proX* | 1 | 35,4 |
| -3,47 | D3NV01 | AZL_015920 | Corrinoid adenosyltransferase | *btuR* | 3 | 22,4 |
| -2,46 | D3P311 | AZL_b05180 | Aspartate ammonia-lyase | *aspA* | 2 | 50,6 |
| -2,29 | D3P1D5 | AZL_a09330 | Amidohydrolase |  | 1 | 35,3 |
| -2,14 | D3NVN0 | AZL_018210 | Extracytoplasmic solute receptor protein |  | 20 | 40,2 |
| -1,93 | D3NT54 | AZL_009450 | C4-dicarboxylate-binding periplasmic protein |  | 16 | 37,5 |
| -1,93 | D3P7I2 | AZL_e02660 | Uncharacterized protein |  | 16 | 24,7 |
| -1,93 | D3NZL4 | AZL_a03120 | Histidine kinase | *cheA* | 1 | 80,3 |
| -1,78 | D3P4Q5 | AZL_c03410 | Uncharacterized protein |  | 4 | 27,6 |
| -1,74 | D3P791 | AZL_e01750 | Cytochrome C signal peptide protein |  | 1 | 16,7 |
| -1,7 | D3P5Y4 | AZL_d02370 | Uncharacterized protein |  | 23 | 41,6 |
| -1,69 | D3P8B9 | AZL_f01380 | dTDP-4-dehydrorhamnose epimerase |  | 4 | 15,8 |
| -1,67 | D3NQP1 | AZL_000820 | SRPBCC family protein |  | 3 | 17,8 |
| -1,52 | D3P2X7 | AZL_b04840 | Sorbitol/mannitol transport system substrate-binding protein | *mtlE* | 11 | 47,3 |
| -1,49 | D3NZK6 | AZL_a03040 | ABC transporter periplasmic substrate-binding protein |  | 15 | 40,1 |
| -1,47 | D3NSQ0 | AZL_007910 | Iron complex outer membrane receptor protein |  | 37 | 74,4 |
| -1,46 | D3NTV4 | AZL_011950 | Secreted protease |  | 16 | 54,1 |
| -1,45 | D3P4Q2 | AZL_c03380 | Fatty-acyl-CoA synthase |  | 2 | 51,2 |
| -1,44 | D3P718 | AZL_e01020 | Flagellin | *fliC* | 16 | 43,4 |
| -1,44 | D3NUV6 | AZL_015470 | Molybdopterin-guanine dinucleotide biosynthesis protein B | *mobB* | 3 | 17,9 |
| -1,43 | D3NW48 | AZL_019890 | Uncharacterized protein |  | 3 | 18,5 |
| -1,42 | D3P162 | AZL_a08600 | Uncharacterized protein |  | 10 | 36,5 |
| -1,42 | D3NZZ0 | AZL_a04380 | Cytochrome C | *cyc* | 3 | 12,9 |
| -1,41 | D3P6R0 | AZL_d05130 | Beta_helix domain-containing protein |  | 33 | 67,9 |
| -1,41 | D3P3Y3 | AZL_c00690 | Putrescine-binding periplasmic protein | *potD* | 15 | 39,3 |
| -1,39 | D3P5V7 | AZL_d02100 | Uncharacterized protein |  | 15 | 24,6 |
| -1,39 | D3NTW6 | AZL_012070 | Flagellar basal-body rod protein FlgC | *flgC* | 7 | 15,2 |
| -1,36 | D3P329 | AZL_b05360 | Uncharacterized protein |  | 18 | 19,1 |
| -1,35 | D3NRT6 | AZL_004770 | Zinc/manganese transport system substrate-binding protein |  | 11 | 32,9 |
| -1,34 | D3P5Y7 | AZL_d02400 | Beta_helix domain-containing protein |  | 55 | 110,9 |
| -1,32 | D3P7A5 | AZL_e01890 | Uncharacterized protein |  | 8 | 31,7 |
| -1,31 | D3NVD7 | AZL_017280 | Uncharacterized protein |  | 13 | 29,5 |
| -1,3 | D3NTD1 | AZL_010220 | 50S ribosomal protein L32 | *rpmF* | 5 | 6,9 |
| -1,3 | D3P6E8 | AZL_d04010 | Single Cache domain-containing protein |  | 10 | 17,3 |
| -1,29 | D3P4Z8 | AZL_c04340 | Uncharacterized protein |  | 26 | 36,6 |
| -1,28 | D3P3T4 | AZL_c00200 | Extracytoplasmic solute receptor protein |  | 6 | 40,8 |
| -1,27 | D3P3X9 | AZL_c00650 | Betaine-homocysteine S-methyltransferase | *bhmT* | 7 | 32,3 |
| -1,24 | D3P3T6 | AZL_c00220 | Spermidine/putrescine transport system substrate-binding protein | *potD* | 15 | 37,7 |
| -1,23 | D3NS18 | AZL_005590 | Branched-chain amino acid transport system substrate-binding protein |  | 11 | 44,9 |
| -1,21 | D3P7D5 | AZL_e02190 | TRAP dicarboxylate transporter |  | 3 | 36,8 |
| -1,18 | D3NX05 | AZL_022960 | Uncharacterized protein |  | 21 | 36,8 |
| -1,18 | D3P0P1 | AZL_a06890 | Endoglucanase |  | 40 | 87,6 |
| -1,17 | D3P6N3 | AZL_d04860 | Caspase family p20 domain-containing protein |  | 29 | 181,7 |
| -1,16 | D3NW01 | AZL_019420 | Cobalamin synthesis protein |  | 10 | 43,7 |
| -1,15 | D3P4G7 | AZL_c02530 | Imelysin-like domain-containing protein |  | 13 | 45 |
| -1,14 | D3NYP8 | AZL_028890 | Protein-export protein SecB | *secB* | 6 | 17,9 |
| -1,14 | D3NZH1 | AZL_a02690 | Poly(3-hydroxybutyrate) depolymerase |  | 9 | 38 |
| -1,13 | D3NUM6 | AZL_014670 | Uncharacterized protein |  | 16 | 41,4 |
| -1,13 | D3P7Z0 | AZL_f00090 | Calcium-binding protein |  | 34 | 83,6 |
| -1,12 | D3NVB2 | AZL_017030 | Sel1 repeat family protein |  | 2 | 37,4 |
| -1,11 | D3NXE5 | AZL_024360 | TRAP dicarboxylate transporter |  | 21 | 36,8 |
| -1,1 | D3P7A4 | AZL_e01880 | Uncharacterized protein |  | 3 | 17,9 |
| -1,1 | D3NVP3 | AZL_018340 | Flagellar basal-body rod protein FlgG | *flgG* | 11 | 27,4 |
| -1,09 | D3P7A6 | AZL_e01900 | Uncharacterized protein |  | 10 | 32,4 |
| -1,06 | D3P338 | AZL_b05450 | Lysozyme |  | 8 | 18,2 |
| -1,05 | D3NT80 | AZL_009710 | Glyceraldehyde-3-phosphate dehydrogenase | *gapA* | 7 | 36,1 |
| -1,04 | D3NXL1 | AZL_025020 | Dihydrolipoyl dehydrogenase | *pdhD* | 30 | 48,7 |
| -1,02 | D3P0A2 | AZL_a05500 | Flagellar hook-associated protein 1 | *flgK* | 32 | 50,4 |
| -1 | D3NYT3 | AZL_a00310 | 50S ribosomal protein L13 | *rplM* | 15 | 17,4 |
| 1,01 | D3NX22 | AZL_023130 | Acetyl-CoA C-acetyltransferase | *atoB* | 28 | 40,3 |
| 1,01 | D3P3R4 | AZL_b01400 | 3-oxoacyl-[acyl-carrier-protein] reductase |  | 2 | 29 |
| 1,01 | D3P891 | AZL_f01100 | Glycosyltransferase |  | 3 | 42,5 |
| 1,02 | D3NYG4 | AZL_028050 | Extracellular ligand-binding receptor |  | 13 | 41,8 |
| 1,04 | D3NYT5 | AZL_a00330 | N-acetyl-gamma-glutamyl-phosphate reductase | *argC* | 15 | 37,8 |
| 1,04 | D3P1Y4 | AZL_b01410 | Tropine dehydrogenase |  | 1 | 27 |
| 1,05 | D3P1R6 | AZL_a10640 | Efflux transporter RND family, MFP subunit |  | 7 | 33,1 |
| 1,06 | D3NSF5 | AZL_006960 | Two-component hybrid sensor and regulator |  | 1 | 72,9 |
| 1,06 | D3P5E5 | AZL_d00480 | 3-carboxy-cis,cis-muconate cycloisomerase | *pcaB* | 2 | 49 |
| 1,08 | D3NSN6 | AZL_007770 | FMN dependent NADH:quinone oxidoreductase | *acpD* | 11 | 21,7 |
| 1,08 | D3NZ10 | AZL_a01080 | 2OG-Fe(II) oxygenase family protein |  | 4 | 39 |
| 1,09 | D3NSL1 | AZL_007520 | Dipeptide epimerase |  | 1 | 34,8 |
| 1,14 | D3NT94 | AZL_009850 | Peptidoglycan-associated lipoprotein | *pal* | 10 | 16,6 |
| 1,16 | D3NU95 | AZL_013360 | proton-translocating NAD(P)(+) transhydrogenase | *pntA* | 3 | 14,4 |
| 1,21 | D3NZH2 | AZL_a02700 | Enolase | *eno* | 1 | 44,7 |
| 1,21 | D3NTY2 | AZL_012230 | ACT domain-containing protein |  | 3 | 20,6 |
| 1,21 | D3NXU8 | AZL_025890 | NADH dehydrogenase | *ndh* | 3 | 47,2 |
| 1,21 | D3P211 | AZL_b01680 | Acyl-CoA oxidase/dehydrogenase |  | 4 | 46 |
| 1,24 | D3NYH4 | AZL_028150 | Methyl-accepting chemotaxis protein |  | 1 | 59,8 |
| 1,24 | D3P260 | AZL_b02170 | 3-hydroxyacyl-CoA dehydrogenase | *fadB* | 4 | 81,1 |
| 1,27 | D3NSG3 | AZL_007040 | DUF3035 domain-containing protein |  | 7 | 21,2 |
| 1,28 | D3NTU9 | AZL_011900 | Resolvase/invertase-type recombinase catalytic domain-containing protein |  | 3 | 22,7 |
| 1,31 | D3NYH9 | AZL_028200 | Long-chain fatty acid transport protein |  | 9 | 43,9 |
| 1,32 | D3P5E3 | AZL_d00460 | Protocatechuate 3,4-dioxygenase, alpha subunit | *pcaG* | 1 | 19,9 |
| 1,35 | D3NTC8 | AZL_010190 | SmpA/OmlA domain protein |  | 4 | 19 |
| 1,43 | D3NRK3 | AZL_003940 | Phage shock protein B |  | 3 | 8,8 |
| 1,47 | D3P192 | AZL_a08900 | Transcriptional regulator |  | 2 | 34,5 |
| 1,48 | D3NV49 | AZL_016400 | Lipoprotein |  | 4 | 23,3 |
| 1,52 | D3P6C8 | AZL_d03810 | Uncharacterized protein |  | 1 | 26,6 |
| 1,54 | D3P297 | AZL_b02540 | Sulfonate monooxygenase |  | 1 | 40,8 |
| 1,6 | D3P289 | AZL_b02460 | Threonine synthase | *thrC* | 8 | 45,5 |
| 1,6 | D3P259 | AZL_b02160 | Acetyl-CoA acyltransferase | *fadA* | 4 | 39,3 |
| 1,64 | D3NWJ8 | AZL_021390 | NADH-quinone oxidoreductase subunit J | *nuoJ* | 1 | 21,7 |
| 1,67 | D3NSJ7 | AZL_007380 | ATP synthase subunit b | *atpF* | 9 | 22,5 |
| 1,68 | D3NV74 | AZL_016650 | Two-component response regulator modulated CheB methylesterase |  | 1 | 38,3 |
| 1,76 | D3P256 | AZL_b02130 | Enoyl-CoA hydratase | *paaG* | 11 | 28,9 |
| 1,77 | D3NWS9 | AZL_022200 | UDP-glucose 4-epimerase |  | 1 | 33,8 |
| 1,79 | D3P253 | AZL_b02100 | Acyl-CoA dehydrogenase-like |  | 7 | 44,5 |
| 1,86 | D3P252 | AZL_b02090 | Acyl-CoA dehydrogenase |  | 9 | 37,9 |
| 1,87 | D3P1Z2 | AZL_b01490 | Dioxygenase subunit alpha |  | 11 | 50,4 |
| 1,89 | D3P258 | AZL_b02150 | 2-nitropropane dioxygenase |  | 11 | 35,5 |
| 1,91 | D3P2L2 | AZL_b03690 | Aldehyde dehydrogenase | *vdh* | 11 | 50,6 |
| 1,98 | D3NX92 | AZL_023830 | DUF3971 domain-containing protein |  | 2 | 124,6 |
| 2,03 | D3NWP9 | AZL_021900 | Two-component response regulator |  | 2 | 25,8 |
| 2,05 | D3P190 | AZL_a08880 | Uncharacterized protein |  | 8 | 39,9 |
| 2,2 | D3NW70 | AZL_020110 | 2Fe-2S ferredoxin |  | 1 | 11,9 |
| 2,32 | D3P2F2 | AZL_b03090 | Enolase | *eno* | 1 | 44,7 |
| 2,34 | D3P1Z0 | AZL_b01470 | Pyruvate dehydrogenase E1 component, beta subunit | *pdhB* | 5 | 35 |
| 2,4 | D3P5E4 | AZL_d00470 | Protocatechuate 3,4-dioxygenase, beta subunit | *pcaH* | 3 | 27,2 |
| 2,47 | D3P213 | AZL_b01700 | Universal stress protein UspA-like nucleotide-binding protein |  | 3 | 29,2 |
| 2,69 | D3P204 | AZL_b01610 | Enoyl-CoA hydratase | *paaG* | 9 | 32,6 |
| 2,73 | D3NQX3 | AZL_001640 | PrcB C-terminal domain-containing protein |  | 1 | 15,9 |
| 2,76 | D3P216 | AZL_b01730 | Electron transfer flavoprotein beta subunit | *etfB* | 8 | 26,4 |
| 2,76 | D3NVD9 | AZL_017300 | YbjN domain-containing protein |  | 1 | 18,7 |
| 2,97 | D3NS72 | AZL_006130 | histidine kinase |  | 1 | 58 |
| 3,36 | D3P205 | AZL_b01620 | Enoyl-CoA hydratase | *paaG* | 15 | 29,5 |
| 3,39 | D3P247 | AZL_b02040 | Thiolase |  | 4 | 40,3 |
| 3,43 | D3P246 | AZL_b02030 | Acetyl-CoA C-acetyltransferase | *atoB* | 7 | 41,3 |
| 3,48 | D3P188 | AZL_a08860 | Aromatic-ring-hydroxylating dioxygenase, alpha subunit |  | 12 | 47,8 |
| 4,1 | D3P206 | AZL_b01630 | Enoyl-CoA hydratase |  | 16 | 33,4 |
| 4,18 | D3P251 | AZL_b02080 | 3-oxoacyl-[acyl-carrier-protein] reductase | *fabG* | 11 | 26,9 |
| 6,64 | D3P215 | AZL_b01720 | Electron transfer flavoprotein, alpha subunit | *etfA* | 4 | 31,4 |
| 6,64 | D3P254 | AZL_b02110 | CaiB/BaiF family protein |  | 7 | 45,1 |
| 6,64 | D3P7B3 | AZL_e01970 | ABC-type transport auxiliary lipoprotein component domain-containing protein |  | 2 | 22,8 |
| 6,64 | D3P2L1 | AZL_b03680 | Enoyl-CoA hydratase | *ech* | 2 | 31,5 |
| 6,64 | D3P3J1 | AZL_b00670 | Pyruvate dehydrogenase E2 component | *pdhC* | 4 | 27,7 |
| 6,64 | D3P210 | AZL_b01670 | Isovaleryl-CoA dehydrogenase | *ivd* | 3 | 40,1 |
| 6,64 | D3NXB2 | AZL_024030 | tRNA pseudouridine synthase A | *truA* | 1 | 27,2 |
| 6,64 | D3NV47 | AZL_016380 | Isochorismatase domain-containing protein 2A |  | 2 | 19,4 |

**Figure 2:**
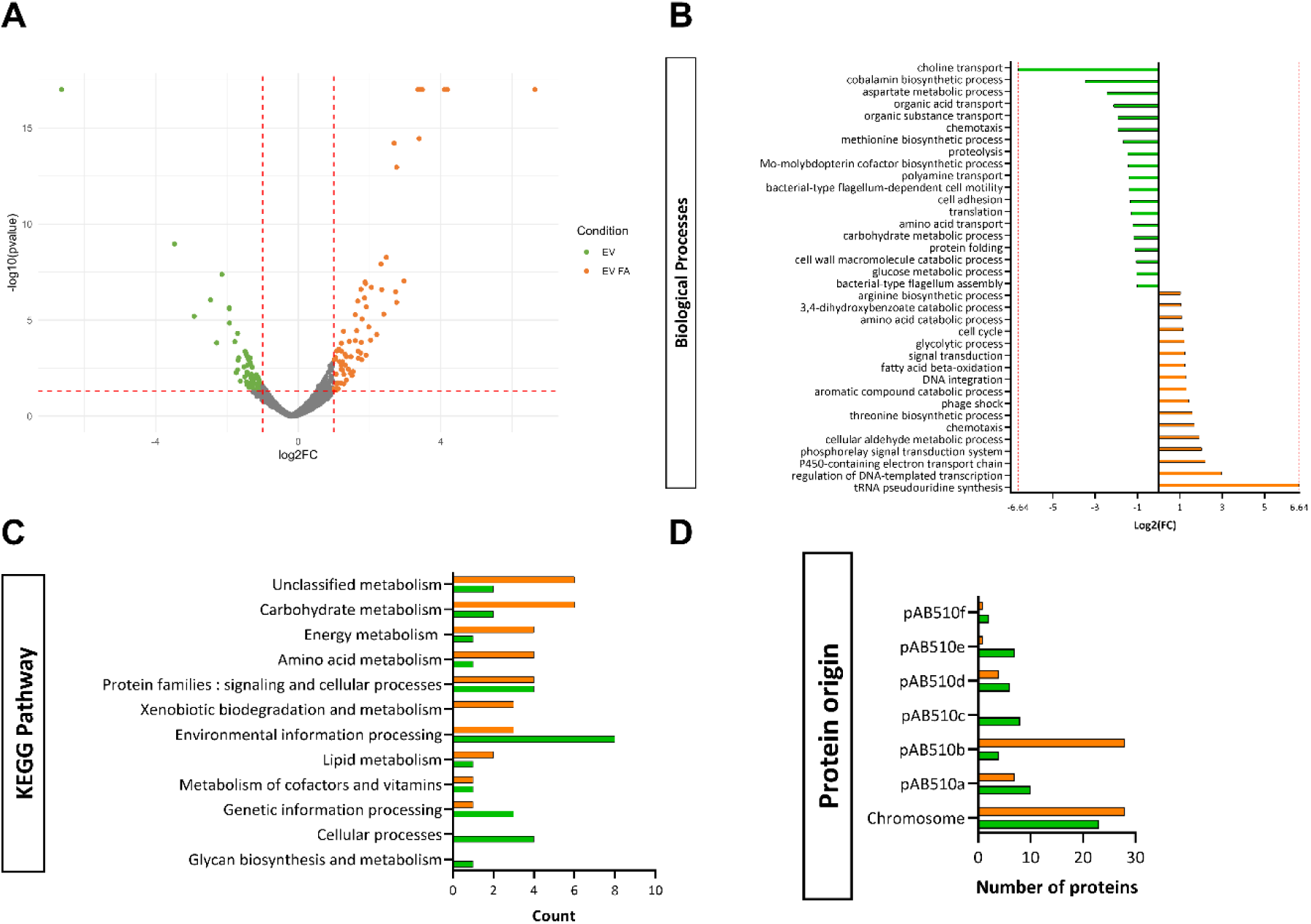
The presence of FA in the growth medium of *Azospirillum* sp. B510 impacts the proteinaceous cargo of EVs. **(A)** Volcano plot representing the modulation of protein accumulation in *Azospirillum* sp. B510 EVs in presence of FA from 3 biological replicates. Colored proteins represent the statistically depleted (green) or enriched (orange) protein (Student’s *t-test*, FC < 0.5 or FC > 2 and *pv* < 0.05, Benjamini-Hochberg correction). **(B)** Biological Process GO categories and **(C)** KEGG Pathway of the *Azospirillum* sp. B510 EVs proteins impacted by the presence of FA. Enrichment is represented as the log2FC of proteins assigned to each GO category. **(D)** Origin of the *Azospirillum* sp. B510 EVs proteins impacted by the presence of FA. EV (green): *Azospirillum* sp. B510 EVs extracted from the ABS condition. EV FA (orange): *Azospirillum* sp. B510 EVs extracted from ABS supplemented with the ABS+FA condition.

#### 3.3.2 Effect of ferulic acid on the lipidic and metabolite content of *Azospirillum* sp. B510 EVs

EVs lipidic membranes are the main structural component of bacterial EVs and dictate their ability to interact with other living cells. Here, we used targeted GC-MS and LC-MS/MS analysis to study the effect of FA on *Azospirillum* sp. B510 EVs lipidic composition. The fatty acid profiles obtained through GC-MS analysis showed that *Azospirillum* sp. B510 EVs were composed of the mono-unsaturated fatty acids. Total phospholipid content analysis by GC-MS revealed that the hydrophilic moieties were mostly linked to mono-unsaturated fatty acid. Further LC-MS/MS analysis on phospholipid content revealed that *Azospirillum* sp. B510 EVs are mainly composed of phosphatidylethanolamine and also include phosphatidylcholine and phosphatidylglycerol (**Table 3**, **Supp. Table 2**).

**Table 2:** Comparison of fatty acid and phospholipid content between *Azospirillum* sp. B510 EV grown with or without FA. Total fatty acid content was analyzed by GC-MS and total lipid content was analyzed by LC-MS/MS. The percentage of each fatty acid or phospholipid is relative to the total fatty acid or phospholipid respectively. PE: Phosphatidylethanolamine ; PC: Phosphatidylcholine; PG: Phosphatidylglycerol. Values represent the mean of three biological replicates.

| Total fatty acid (%) | EV | EV FA |
| --- | --- | --- |
| 16 : 0 | 18.14 | 16.71 |
| 18 : 0 | 19.74 | 17.81 |
| 16 : 1n-7 | 5.8 | 5.41 |
| 18 : 1n-7 | 49.44 | 53.52 |
| 18 : 1n-9c | 6.88 | 6.56 |
| **Total phospholipid (%)** |  |  |
| 16 : 0 | 9.94 | 11.66 |
| 18 : 0 | 6.25 | 10.80 |
| 16 : 1n-7 | 9.88 | 8.19 |
| 18 : 1n-7 | 65.68 | 62.16 |
| 18 : 1n-9c | 5.26 | 7.19 |
| **Phospholipid type (%)** |  |  |
| PE | 80.99 | 80.65 |
| PC | 12.96 | 13.42 |
| PG | 6.05 | 5.93 |

**Table 3:**
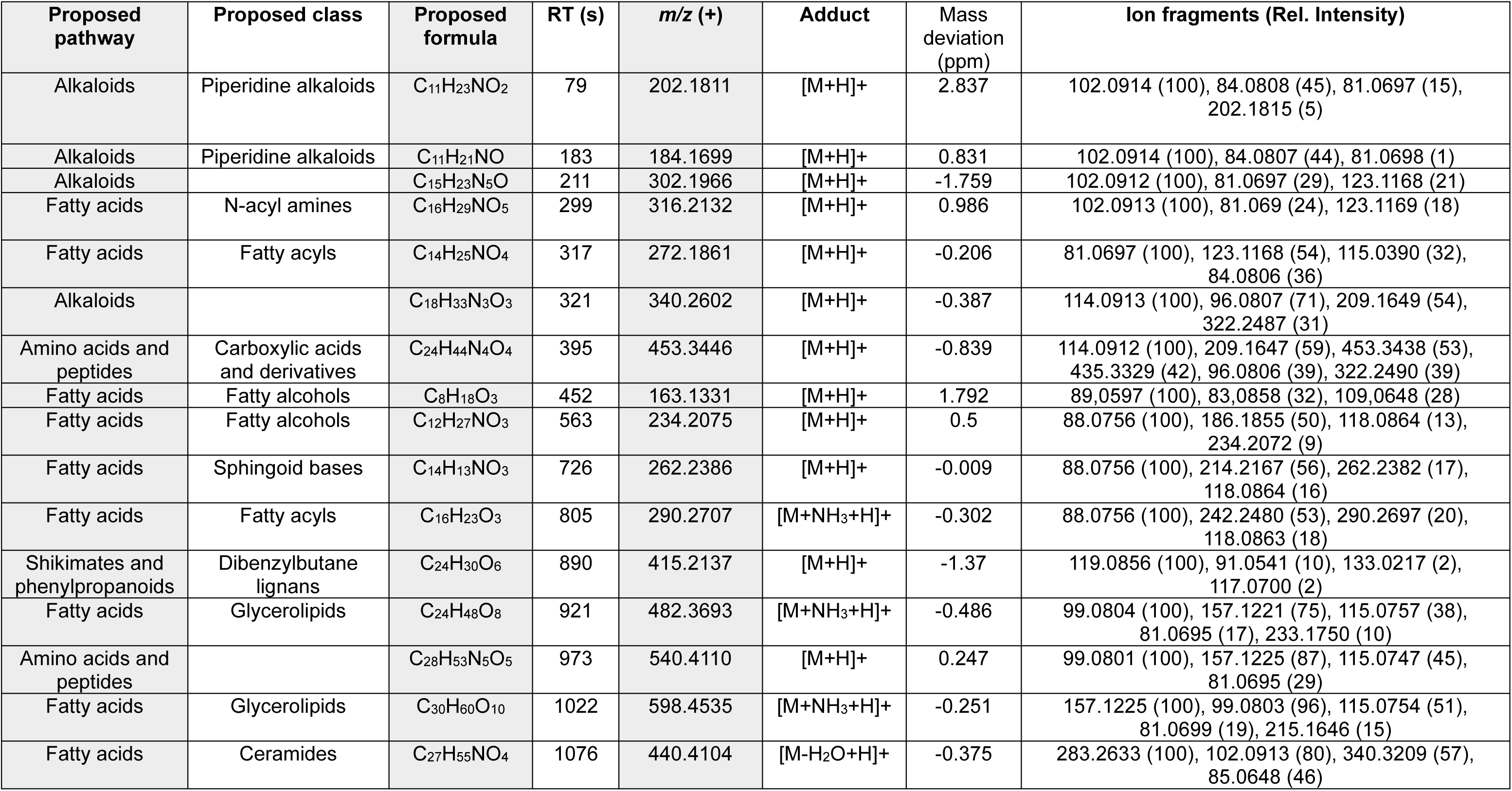
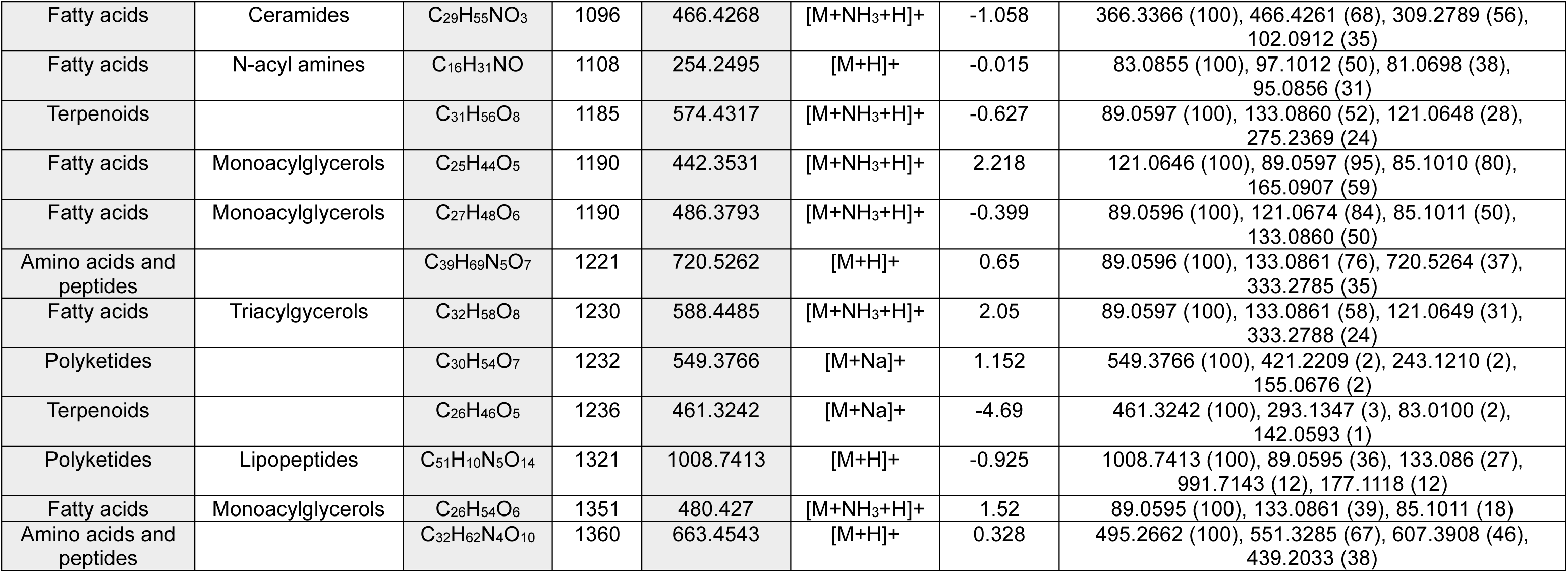
UHPLC-DAD-ESI-MS data of detected and annotated metabolites in *Azospirillum* sp. B510 EVs. Proposed pathways and proposed classes were obtained using the SIRIUS’s CANOPUS module. Compounds were annotated when at least present in 2 of the 3 biological EV and EV FA replicates.

The untargeted metabolomic analysis of EVs from *Azospirillum* sp. B510 grown with or without FA using UHPLC-UV/DAD-ESI-MS-QTOF allowed the annotation of 28 of the most abundant metabolites in EVs samples (**Table 2**). Using CANOPUS, annotated metabolites were mostly assigned to the proposed pathway fatty acids, amino acids and peptides. Iodixanol solution is the main gradient density solution used in EV studies for their purification. Most studies report that washing with dH_2_O (or buffer) is sufficient to remove iodixanol from EV samples. In our case, despite the wash of EVs samples following GDUC, iodixanol ions (*m/z* (+) = 1550.7250) were found at the same intensity in both EV and EV FA samples (**Supp. Fig. 7**).

Every compound detected in EV samples was also present in EV FA samples, indicating that FA did not impact the metabolite and lipid content of *Azospirillum* sp. B510 EVs in the present analysis.

### 3.4 Effect of Azospirillum sp. B510 EVs on Solanum lycopersicum

To study the impact of the cargoes of *Azospirillum* sp. B510 EVs, young tomato plants (14 days of growth) were inoculated on the root with 1.10^9^ EVs particles from EV or EV FA condition. This quantity of EVs was determined as the quantity purified from 5.10^8^ *Azospirillum* sp. B510 cells. In *Azospirillum*-plant interactions studies, this number is classically used as an inoculum (Valette *et al*., 2020). In order to test the persistence and possible systemic effects of EVs on *S. lycopersicum*, tomato roots and aerial tissues were harvested either 18h or 5 days after inoculation.

#### 3.4.1 Effect of *Azospirillum* sp. B510 EVs on the amino acid content of *S. lycopersicum*

As amino acids play a crucial part in plant-microbe interactions and are the backbones of specialized metabolites, we analysed the concentration of 24 amino acids in both roots and aerial parts tissues using HPLC-UV/DAD-FLD. The application of both types of EV on *S. lycopersicum* roots did not show any significant effect on the concentration of the 24 tested amino acids in root tissues compared to the Control condition (data not shown). However, in aerial parts tissues, the concentration of 11 amino acids out of 24 was significantly impacted 18 hpi : α-aminobutyric acid, arginine, lysine and isoleucine were under-abundant in both EV and EV FA conditions compared to the Control condition. Phenylalanine, serine and threonine were under-abundant only in the EV condition compared to the Control condition. Conversely, asparagine and histidine were under-abundant only in the EV FA condition compared to the Control condition. Aspartic acid was under-abundant in the EV FA condition compared to the EV condition but not to the Control condition. Ornithine was the only amino acid to be over-abundant 18 hpi, and solely in the EV condition (**Fig. 3 A**). Only 5 amino acids were significantly impacted in their concentration at 5 dpi. Citrulline and glycine were over-abundant in both EV conditions and EV FA condition compared to the Control condition respectively. Ornithine was still significantly over-abundant in EV condition compared to EV FA condition but not to the Control condition. Proline was under-abundant in EV condition compared to the Control condition and tyrosine was under-abundant in EV condition compared to the EV FA condition (**Fig. 3 B**). Overall, *Azospirillum* sp. B510 EVs induced a systemic modification of amino acids profiles of *S. lycopersicum*. These modifications were shown to depend on both the EVs type and application time.

**Figure 3:**
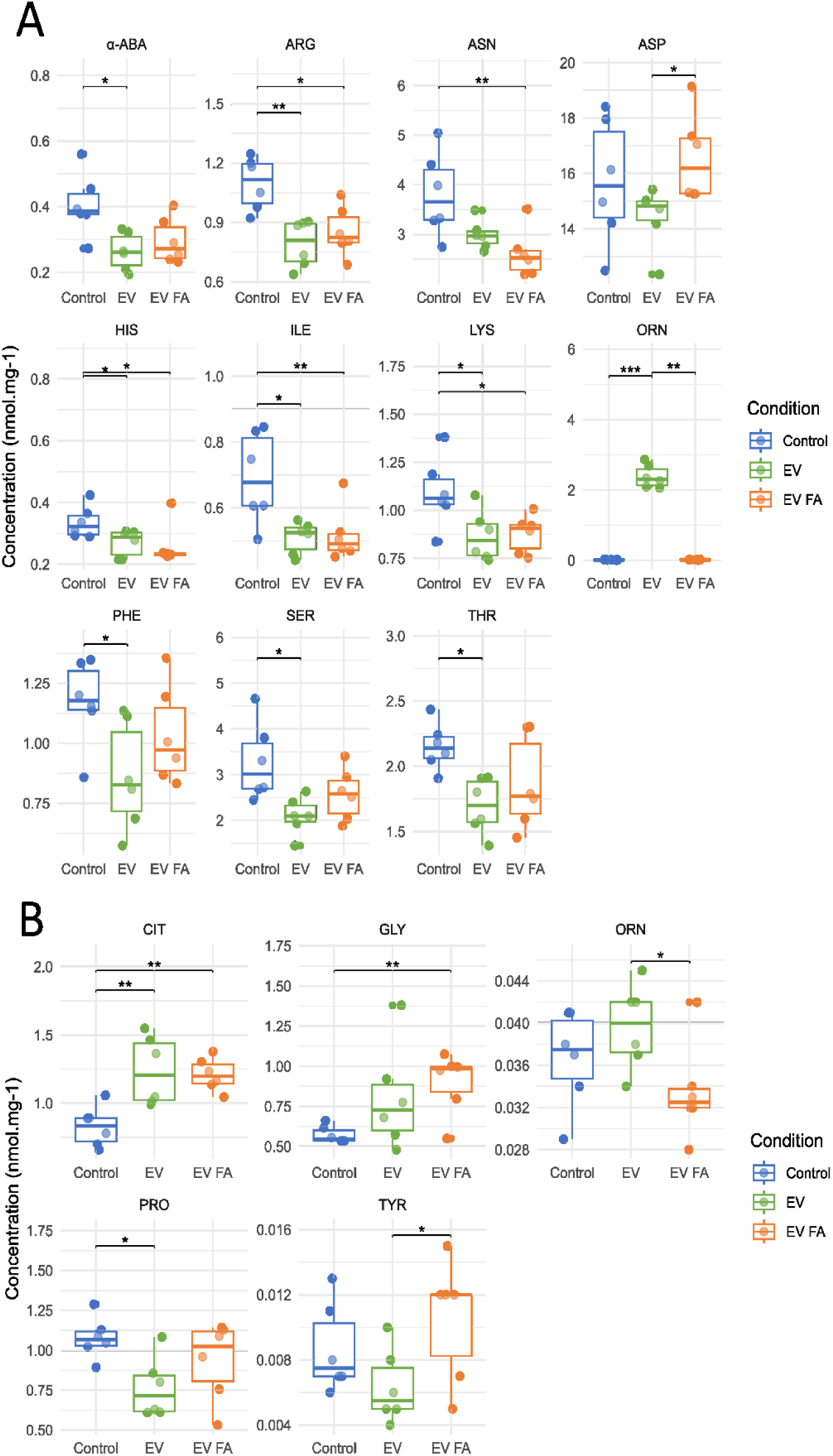
Impact of *Azospirillum* sp. B510 EVs on the amino acid content of *S. lycopersicum* aerial parts. Amino acid concentration was determined using HPLC-UV/DAD-FLD after inoculation of EV, EV FA or water (Control) conditions at the roots of *S. lycopersicum.* Amino acids were quantified 18 hpi **(A)** or 5 dpi **(B)**. α-ABA: α-aminobutyric acid; ARG: arginine; ASN: asparagine; ASP: aspartic acid; CIT: citrulline; GLY: glycine; HIS: histidine ; ILE: isoleucine; LYS: lysine; ORN: ornithine; PHE: phenylalanine; PRO: proline; SER: serine; THR: threonine; TYR: tyrosine. Results show the mean concentration (nmol.mg^−1^) of each amino acid (n = 6, 10 plants/pool, Tukey analysis, * *pv* > 0.05, ** *pv* > 0.01).

#### 3.4.2 Effect of *Azospirillum* sp. B510 EVs on the specialized metabolome of *S. lycopersicum*

We investigated whether the application of *Azospirillum* sp. B510 EVs on *S. lycopersicum* roots had an effect on the tomato specialized metabolome. We thus compared metabolites contents in roots and aerial parts at 18 hpi or 5 dpi of EV or EV FA, to the uninoculated roots (i.e. water Control condition).

Analysis of the data obtained by UHPLC-ESI-MS-QTOF led to the obtention of 4 data matrices corresponding to metabolites profiles of roots and aerial parts 18 hpi and 5 dpi. As iodixanol ions were found in the analysis of plant metabolites, a control experiment was set up to make sure that iodixanol alone would not induce a metabolic response in *S. lycopersicum*. Using the same method, 60 tomato plants were inoculated with 0.01% iodixanol (iodixanol quantity found in *Azospirillum* sp. B510 EVs metabolite analysis, see **3.3.2**) and compared to 60 plants from the Control condition (water). After 18 hpi, no effect of iodixanol on plant metabolites was observed. Thus, iodixanol ions were subsequently removed from tomato root metabolites profiles matrices.

PLS-DA analysis performed on the roots metabolites 18 hpi matrix showed a clear separation between the EV and the Control condition and to a lesser extent, a separation between EV FA and Control condition (**Fig. 4 A**). It also allowed to highlight the most discriminant ions between those conditions (4202 ions, VIP score > 1). PLS-DA analysis was not applicable on the matrix obtained 5 dpi (Q2Y score < 0.1), indicating that both EV and EV FA did not induce any noticeable effect on *S. lycopersicum* root specialized metabolites profiles 5 dpi. On aerial parts, PLS-DA analysis showed a clear separation between the 3 tested conditions 18 hpi and this separation was still observable at 5 dpi (**Fig. 4 B, C**). It allowed the highlight of 4075 and 4071 discriminant ions respectively (VIP score > 1). In the 3 cases where PLS-DA analysis was done, the application of EV and EV FA seemed to reduce the variability of specialized plant metabolites profiles.

**Figure 4:**
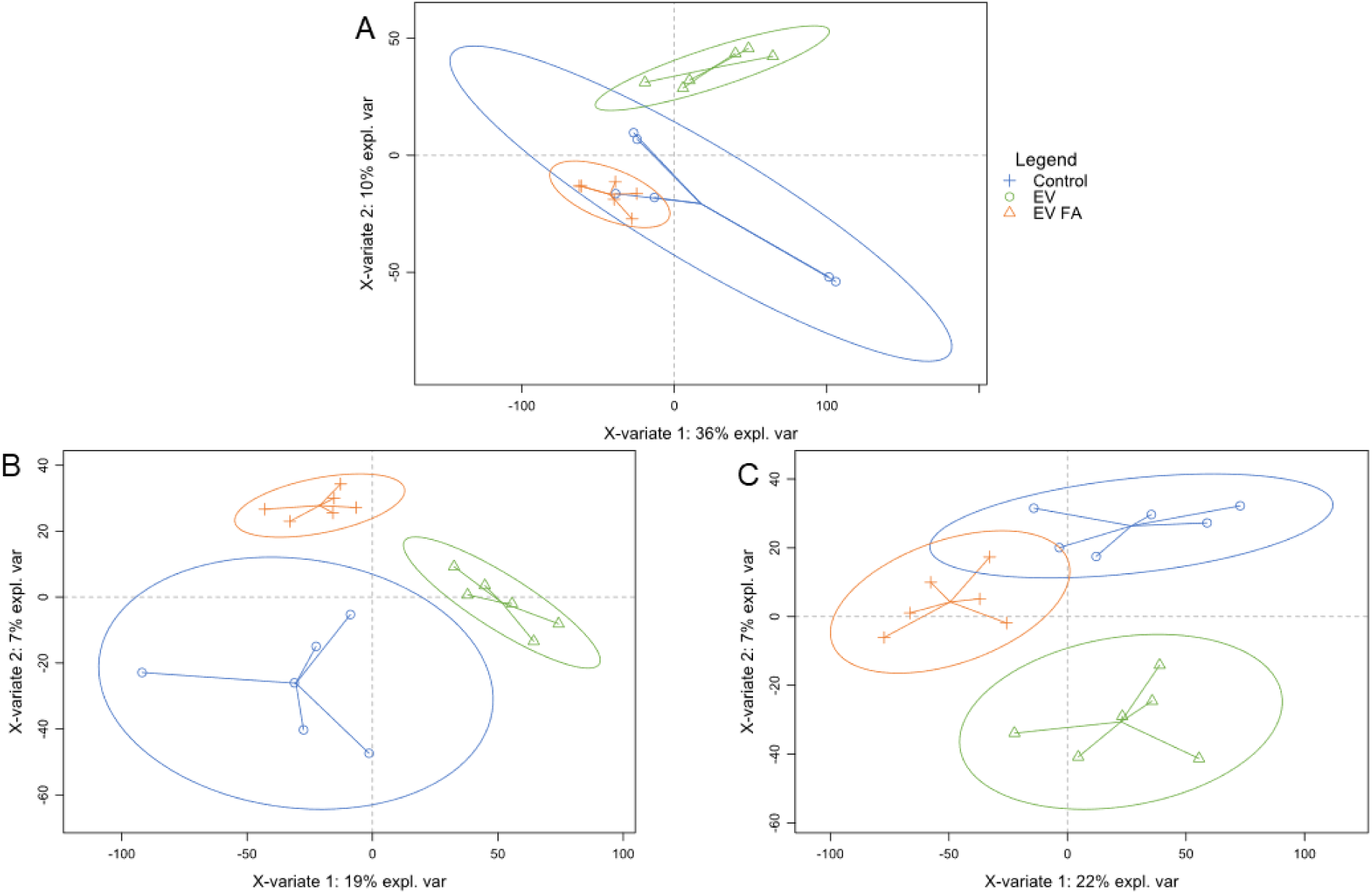
Comparison of *S. lycopersicum* specialized metabolites profiles following the application of *Azospirillum* sp. B510 EVs. PLS-DA analyses were performed on data obtained after the inoculation of either EV, EV FA or water (Control condition). Each point represents the extract of pooled samples of the same treatment and harvest time (10 plants/pool). **(A)** Metabolite profiles of root extracts at 18 hpi. **(B)** Metabolite profiles of aerial parts extracts at 18 hpi. **(C)** Metabolite profiles of aerial parts extracts at 5 dpi.

#### 3.4.3 Annotation of discriminant compounds

Annotation of discriminating compounds (VIP score > 1) in each plant compartment was carried out on QC samples using SIRIUS on MS/MS spectra and the resulting proposed formulas were manually compared to the COCONUT and KNApSAcK databases and to the literature when available. The CANOPUS module of SIRIUS allowed the categorization of molecules into putative molecular classes. Each table summarizes the putative class, putative identity, most probable formula, *m/z* and VIP score of discriminating ions for which an annotation was obtained in extracts from tomato roots (**Table 4**) or aerial parts (**Table 5**). In the root parts, the most represented metabolite class amongst the annotated ions was HCAs and derivatives. Indeed, 8 discriminating ions were annotated as HCAAs: *N*-benzoyl-putrescine, *N*-sinapoylputrescine, *N-*(*p*-coumaroyl)-putrescine, *N*-(*p*-coumaroyl)-spermidine, *N*-feruloylcadaverine, *N*-cis-feruloyloctopamine, *N*-(feruloyl-O-hexoside)-tyramine and *N*-trans-isoferuloyltyramine. The application of EV FA induced a depletion of all those HCAAs in tomato roots compared to the EV and Control condition at 18 hpi (**Supp. Fig. 8**). In the aerial parts, most of the annotated discriminating ions were steroidal alkaloids belonging to the tomatidine family as tomatine isomers, hydroxytomatine isomers, dehydrotomatine isomers and acetoxytomatine isomers. The abundance of most of these molecules was dependent on the type of EVs and the contact time. These steroidal alkaloids were more abundant in the EV than in the EV FA or Control condition 18 hpi while they were more abundant in the EV FA than in EV or Control condition 5 dpi (**Supp. Fig. 9**). Moreover, two annotated HCAAs in the aerial parts, the *N*-feruloylputrescine and the *N*-feruloyltyramine, were shown to be under-abundant in the EV FA condition 18 hpi compared to the EV and Control condition. However, these compounds seemed to be more abundant in EV FA at 5 dpi. Overall, *Azospirillum* sp. B510 EVs induced a local and systemic modification of specialized metabolites profiles of *S. lycopersicum*. These modifications were shown to depend on both the EVs type and application time.

**Table 4:** UHPLC-DAD-ESI-MS data of detected and annotated VIP metabolites in *Solanum lycopersicum* roots. Proposed pathways and proposed classes were obtained using the SIRIUS’s CANOPUS module (a). When available, mass spectra were compared to standard molecules (b) or to the following literature: (c) Itkin et *al.*, 2011, (d) Li et *al.*, 2018, (e) Dastmalchi et al., 2014, (f) Nohara et *al.*, 2010.

| Proposed pathway | Proposed class | Putative metabolite | Proposed formula | RT (s) | m/z (+) | Adduct | Mass deviation (ppm) | VIP Score | Ref |
| --- | --- | --- | --- | --- | --- | --- | --- | --- | --- |
| Alkaloids | Piperidine alkaloids | O-Acetylgeniposidic acid | C<sub>18</sub>H<sub>24</sub>O<sub>11</sub> | 198 | 186.1853 | [M+Na]<sup>+</sup> | 0.007 | 1.08 | a |
| Alkaloids | Hydroxycinnamic acid | N-Benzoyl-putrescine | C<sub>11</sub>H<sub>16</sub>N<sub>2</sub>O | 268 | 193.1340 | [M+H]<sup>+</sup> | 0.275 | 1.46 | a |
| Alkaloids | Hydroxycinnamic acid | N-Sinapoyl-putrescine | C<sub>15</sub>H<sub>22</sub>N<sub>2</sub>O<sub>4</sub> | 273 | 295.1652 |  |  | 1.48 | a |
| Phenylpropanoids and derivatives | Hydroxycinnamic acid | N-Coumaroyl-putrescine | C<sub>13</sub>H<sub>18</sub>N<sub>2</sub>O<sub>2</sub> | 275 | 235.1441 | [M+H]<sup>+</sup> | -0.261 | 1.25 | a, e |
| Benzenoids | Benzene and substituted derivatives | 1-Benzoylpiperidine | C<sub>12</sub>H<sub>15</sub>NO | 296 | 190.1227 | [M+H]<sup>+</sup> | -0.832 | 1.34 | a |
| Shykimates and phenylpropanoids | Organooxygen compounds | Zizybesoide I | C<sub>19</sub>H<sub>28</sub>O<sub>11</sub> | 320 | 455.1526 | [M+Na]<sup>+</sup> | -0.103 | 2.02 | a, f |
|  | Hydroxycinnamic acids and derivatives | N-(p-Coumaroyl)-spermidine | C<sub>24</sub>H<sub>44</sub>N<sub>4</sub>O<sub>4</sub> | 324 | 292.2019 |  |  | 1.05 | b |
|  | Hydroxycinnamic acids and derivatives | N-Feruloyl-cadaverine | C<sub>15</sub>H<sub>22</sub>N<sub>2</sub>O<sub>3</sub> | 366 | 279.1704 | [M+H]<sup>+</sup> | 0.148 | 1.32 | a |
| Shykimates and phenylpropanoids | Phenylethanoids | Benzyl β-primeveroside | C<sub>18</sub>H<sub>26</sub>O<sub>10</sub> | 381 | 402.1535 | [M+NH<sub>3</sub>+H]<sup>+</sup> | 1.025 | 1.51 | a |
|  | Indole and derivatives | 2-[1-(4,5-dihydroxy-3-[[3,4,5-trihydroxy-6-(hydroxymethyl)oxan-2- | C<sub>21</sub>H<sub>27</sub>NO<sub>11</sub> | 423 | 469.158 | [M+Na]<sup>+</sup> | -0.152 | 1.59 | a |
|  |  | yl]oxy}oxan-2-yl)-1H-indol-3-yl]acetic acid |  |  |  |  |  |  |  |
| Alkaloids | 1-pyranosylindoles | 1-(1-B-Glucopyranosyl)-1H-Indole-3-Acetic Acid | C<sub>16</sub>H<sub>19</sub>NO<sub>7</sub> | 428 | 337.1163 | [M+Na]<sup>+</sup> | 0.996 | 1.85 | a |
|  | Fatty acyls | 2-[[6-(hex-3-en-1-yloxy)-3,4,5-trihydroxyoxan-2-yl]methoxy]-6-(hydroxymethyl)oxane-3,4,5-triol | C<sub>18</sub>H<sub>32</sub>O<sub>11</sub> | 467 | 447.1842 | [M+Na]<sup>+</sup> | -1.105 | 1.62 | a |
| Shykimates and phenylpropanoids | Phenolic acids | Violutoside | C<sub>19</sub>H<sub>26</sub>O<sub>12</sub> | 472 | 469.1324 | [M+Na]<sup>+</sup> | -1.406 | 1.01 | a |
| Fatty acids | Fatty acyl glycosides of mono- and disaccharides | 4-[methylpentoxy]oxane-3,4,5-triol | C<sub>19</sub>H<sub>36</sub>O<sub>12</sub> | 549 | 479.2100 | [M+Na]<sup>+</sup> | -0.093 | 1.35 | a |
| Shykimates and phenylpropanoids | Hydroxycinnamic acids and derivatives | *N*-Feruloyloctopamine | C<sub>18</sub>H<sub>19</sub>O<sub>5</sub> | 693 | 352.1156 | [M+Na]<sup>+</sup> | -1.678 | 1.48 | a, d |
| Shykimates and phenylpropanoids | Hydroxycinnamic acids and derivatives | N(Feruloyl-O-hexoside)tyramine | C<sub>24</sub>H<sub>29</sub>NO | 710 | 498.1735 | [M+Na]<sup>+</sup> | 0.294 | 1.21 | a |
| Shykimates and phenylpropanoids | Hydroxycinnamic acids and derivatives | Demethyleugenol 4-O-Beta-D-Xylopyranosyl-(1-6)-O-Beta-D-Glucopyranoside | C<sub>20</sub>H<sub>28</sub>O<sub>11</sub> | 741 | 462.1972 | [M+NH<sub>3</sub>+H]<sup>+</sup> | -0.73 | 1.12 | a |
| Terpenoids |  | Phenethyl Rutinoside | C<sub>20</sub>H<sub>30</sub>O<sub>10</sub> | 753 | 448.2183 | [M+NH<sub>3</sub>+H]<sup>+</sup> | 1.708 | 1.23 | a |
| Shykimates and phenylpropanoids | Organic oxygen compounds | 2-[(4,5-dihydroxy-2-[[3-hydroxy-1-methyl-7-(prop-1-en-2-yl)- | C<sub>26</sub>H<sub>42</sub>O<sub>12</sub> | 984 | 564.3015 | [M+NH<sub>3</sub>+H]<sup>+</sup> | 0.0667 | 1.52 | a |
|  |  | 1,2,3,4,5,6,7,8-octahydronaphthalen-2-yl]oxy}-6-(hydroxymethyl)oxan-3-yl]oxy]-6-(hydroxymethyl)oxane-3,4,5-triol |  |  |  |  |  |  |  |
| Shykimates and phenylpropanoids | Hydroxycinnamic acids and derivatives | *N*-trans-isoferuloyltyramine | C<sub>18</sub>H<sub>19</sub>NO<sub>4</sub> | 1035 | 314.1388 | [M+H]<sup>+</sup> | 0.164 | 1.51 | a |
| Fatty acids | Fatty acyl glycosides of mono- and disaccharides | 2-methyl-6-[(3,4,5-trihydroxy-6-{[4-hydroxy-3-(propan-2-yl)pentyl]oxy}oxan-2-yl)methoxy]oxane-3,4,5-triol | C<sub>20</sub>H<sub>38</sub>O<sub>11</sub> | 1056 | 472.2753 | [M+NH<sub>3</sub>+H]<sup>+</sup> | 0.196 | 2.06 | a |
|  | Organic oxygen compounds | 2-(3-heptyl-5-hydroxyphenoxy)-6-(hydroxymethyl)-5-methoxyoxane-3,4-diol | C<sub>20</sub>H<sub>32</sub>O<sub>7</sub> | 1145 | 402.2488 | [M+NH<sub>3</sub>+H]<sup>+</sup> | -0.405 | 1.76 | a |
|  | Fatty acyls | 5-[(diaminomethylidene)amino]-2-{3-methyl-2-[(5-oxopyrrolidin-2-yl)formamido]butanamido}pentanoic acid | C<sub>19</sub>H<sub>536</sub>O<sub>10</sub> | 1160 | 442.2653 | [M+NH<sub>3</sub>+H]<sup>+</sup> | 1.175 | 1.85 | a |
| Steroidal alkaloids | Steroids and steroid derivatives | β-Tomatidine | C<sub>45</sub>H<sub>75</sub>NO<sub>17</sub> | 1233 | 902.5111 | [M+H]<sup>+</sup> | 0.285 | 1.18 | a, c |
| Steroidal alkaloids | Steroids and steroid derivatives | Tomatine | C<sub>50</sub>H<sub>83</sub>NO<sub>21</sub> | 1240 | 1056.535 | [M+Na]<sup>+</sup> |  | 1.01 | a, b |
|  | Fatty acyls | 2-(octyloxy)-6-[(3,4,5-trihydroxyoxan-2-yl)oxy]methyl}oxane-3,4,5-triol | C<sub>19</sub>H<sub>36</sub>O<sub>10</sub> | 1245 | 442.2647 | [M+NH<sub>3</sub>+H]<sup>+</sup> | 0.121 | 1.11 | a |
|  | Prenol lipids | Pisumoside B | C<sub>32</sub>H<sub>52</sub>O<sub>16</sub> | 1342 | 710.3596 | [M+NH<sub>3</sub>+H]<sup>+</sup> | -0.343 | 1.44 | a |

**Table 5:** UHPLC-DAD-ESI-MS data of detected and annotated VIP metabolites in *Solanum lycopersicum* aerial parts. Proposed pathways and proposed classes were obtained using the SIRIUS’s CANOPUS module (a). When available, mass spectra were compared to standard molecules (b) or to the following literature: (c) Itkin et *al.*, 2011, (d) Li et *al.*, 2018, (e) Dastmalchi et al., 2014, (f) Nohara et *al.*, 2010.

| Proposed pathway | Proposed class | Putative metabolite | Proposed formula | RT (s) | m/z (+) | Adduct | Mass deviation (ppm) | VIP Score 18 h | VIP Score 5 d | Ref |
| --- | --- | --- | --- | --- | --- | --- | --- | --- | --- | --- |
| Alkaloids | Simple indole alkaloids | Indole-3-acetaldehyde | C<sub>10</sub>H<sub>9</sub>NO | 206 | 160.0758 | [M+K]<sup>+</sup> | 0.693 | 0.38 | 1.41 | a |
| Amino acids and peptides | Amino acids | L-Phenylalaline | C<sub>9</sub>H<sub>11</sub>NO<sub>2</sub> | 232 | 166.0110 | [M+H]<sup>+</sup> | -0.474 | 0.88 | 1.66 | a |
| Alkaloids | Hydroxycinnamic acid | Caffeoyl-putrescine | C<sub>13</sub>H<sub>18</sub>N<sub>2</sub>O<sub>3</sub> | 279 | 251.1395 | [M+H]<sup>+</sup> | 0.377 | 0.71 | 1.6 | d,e |
| Alkaloids | Hydroxycinnamic acid | Feruloyl-putrescine | C<sub>14</sub>H<sub>20</sub>N<sub>2</sub>O<sub>3</sub> | 275 | 265.1547 | [M+H]<sup>+</sup> | 0.444 | 1.31 | 1.65 | a, e |
| Shikimates and Phenylpropanoids | Organooxygen compounds | Zizybeoside I | C<sub>19</sub>H<sub>28</sub>O<sub>11</sub> | 396 | 455.1522 | [M+Na]<sup>+</sup> | 1.46 | 0.98 | 1.26 | a, f |
| Shikimates and phenylpropanoids | Organooxygen compounds | Icariside F2 | C<sub>18</sub>H<sub>26</sub>O<sub>10</sub> | 466 | 425.1420 | [M+Na]<sup>+</sup> | -0.245 | 1.28 | 1.74 | a |
| Shikimates and Phenylpropanoids | Flavonoids | Sophoraflavonolside | C<sub>27</sub>H<sub>30</sub>O<sub>16</sub> | 691 | 611.1608 | [M+H]<sup>+</sup> | 0.003 | 0.78 | 1.43 | a |
| Steroidal alkaloids | Steroids and steroid derivatives | Hydroxytomatine isomer | C<sub>50</sub>H<sub>83</sub>NO<sub>22</sub> | 863 | 1050.5481 | [M+H]<sup>+</sup> | -0.282 | 2.29 | 1.75 | a, c |
| Steroidal alkaloids | Steroids and steroid derivatives | Dehydrotomatine isomer | C<sub>50</sub>H<sub>81</sub>NO<sub>21</sub> | 902 | 1032.5382 | [M+H]<sup>+</sup> | 0.088 | 1.61 | 2.2 | a, c |
| Steroidal alkaloids | Steroids and steroid derivatives | Hydroxytomatine isomer | C<sub>50</sub>H<sub>83</sub>NO<sub>22</sub> | 972 | 1050.5484 | [M+H]<sup>+</sup> | -0.842 | 2.42 | 0.47 | a, c |
| Steroidal alkaloids | Steroids and steroid derivatives | Hydroxytomatine isomer | C<sub>50</sub>H<sub>83</sub>NO<sub>22</sub> | 1026 | 1050.5486 | [M+H]<sup>+</sup> | 0.677 | 1.32 | 1.93 | a, c |
| Shikimates and Phenylpropanoids | Hydroxycinnamic acids and derivatives | Feruloyl-tyramine | C<sub>18</sub>H<sub>19</sub>NO<sub>4</sub> | 1052 | 314.1387 | [M+H]<sup>+</sup> | -1.085 | 0.33 | 1.04 | e |
| Steroidal alkaloids | Steroids and steroid derivatives | Dehydrotomatidine tetrahexoside isomer | C<sub>51</sub>H<sub>83</sub>NO<sub>22</sub> | 1054 | 1062.5476 | [M+H]<sup>+</sup> | 0.252 | 1.87 | 1.16 | a, c |
| Steroidal alkaloids | Steroids and steroid derivatives | Solasodoside A | C<sub>51</sub>H<sub>82</sub>O<sub>21</sub> | 1057 | 1030.5218 | [M+H]<sup>+</sup> | 0.097 | 1.98 | 1.04 | a |
| Steroidal alkaloids | Steroids and steroid derivatives | Unidentified glycoalkaloid | C<sub>50</sub>H<sub>81</sub>N<sub>3</sub>O<sub>20</sub> | 1082 | 1044.5374 | [M+H]<sup>+</sup> | -0.179 | 0.43 | 1.36 | a, c |
| Steroidal alkaloids | Steroids and steroid derivatives | Dehydrotomatine isomer | C<sub>50</sub>H<sub>81</sub>NO<sub>21</sub> | 1087 | 1032.5389 | [M+H]<sup>+</sup> | -1.548 | 1.81 | 1.12 | a, c |
| Steroidal alkaloids | Steroids and steroid derivatives | Dehydrotomatidine tetrahexoside isomer | C<sub>51</sub>H<sub>83</sub>NO<sub>22</sub> | 1178 | 1062.5481 | [M+H]<sup>+</sup> | 0.827 | 1.51 | 1.38 | a, c |
| Steroidal alkaloids | Steroids and steroid derivatives | Dehydrotomatine isomer | C<sub>50</sub>H<sub>81</sub>NO<sub>21</sub> | 1201 | 1032.5414 | [M+H]<sup>+</sup> | -0.798 | 1.09 | 0.97 | a, c |
| Steroidal alkaloids | Steroids and steroid derivatives | Tomatidine tetrahexoside | C<sub>51</sub>H<sub>85</sub>NO<sub>22</sub> | 1223 | 1064.5645 | [M+H]<sup>+</sup> | -0.859 | 1.36 | 1.73 | a, c |
| Steroidal alkaloids | Steroids and steroid derivatives | Tomatine isomer | C<sub>50</sub>H<sub>83</sub>NO<sub>21</sub> | 1228 | 1034.5530 | [M+H]<sup>+</sup> |  | 0.66 | 1.32 |  |
| Steroidal alkaloids | Steroids and steroid derivatives | Acetoxytomatine isomer | C<sub>52</sub>H<sub>85</sub>NO<sub>23</sub> | 1231 | 1092.5587 | [M+H]<sup>+</sup> | 0.536 | 2.02 | 1.49 | a, c |
| Steroidal alkaloids | Steroids and steroid derivatives | Tomatine | C<sub>50</sub>H<sub>83</sub>NO<sub>21</sub> | 1240 | 1034.5589 | [M+H]<sup>+</sup> |  | 1.31 | 0.1 | b |
| Steroidal alkaloids | Steroids and steroid derivatives | Tomatine isomer | C<sub>50</sub>H<sub>83</sub>NO<sub>21</sub> | 1269 | 1034.5556 | [M+H]<sup>+</sup> |  | 1.15 | 1.48 |  |
| Steroidal alkaloids | Steroids and steroid derivatives | Tomatidine dihexoside dipentoside | C<sub>49</sub>H<sub>81</sub>O<sub>20</sub> | 1294 | 1004.5428 | [M+H]<sup>+</sup> | 1.109 | 1.48 | 1.6 | a, c |

#### 3.4.4 Effect of *Azospirillum* sp. B510 EVs on the expression of defense gene in *S. lycopersicum*

As an effect of *Azospirillum* sp. B510 EVs was observed on the defense molecules in aerial parts of *S. lycopersicum*, we tested the effect of EVs on the expression of tomato defense genes using a biomolecular tool allowing the study of the expression of 28 defense genes (**Supp. Table 3**). At 18 hpi, the *PR14* (lipid transport) and *PPO* (phenylpropanoids pathway) were significatively repressed in *S. lycopersicum* leaves of plants treated with EV FA compared to the Control condition whereas the expression of *CAD* (cell-wall modification), *ANS* (phenylpropanoids pathway) and *TPS* (isoprenoids pathway) were significatively induced in *S. lycopersicum* leaves of plants treated with EV FA. Concerning hormonal signaling, *EIN3* (ethylene pathway) and *JAR* (jasmonic acid pathway) were significatively repressed in *S. lycopersicum* leaves after the application of both EV and EV FA compared to the Control condition. *ACCO* (ethylene pathway) was repressed only in presence of EV FA and *EDS1* (salicylic acid pathway) was over-expressed only in presence of EV FA compared to the other conditions (**Fig. 5 A**). At 5 dpi, *EIN3* was still repressed in both EV and EV FA compared to the Control condition while *HMGR* (isoprenoid pathway) was induced in EV FA compared to the EV condition and *TPS* expression (isoprenoid pathway) was induced in both EV and EV FA compared to the Control condition (**Fig. 5 B**). The expression of *S. lycopersicum* defense gene in leaves was shown to be modulated upon EVs application to the roots. This modulation was different according to the EVs type and application time.

**Figure 5:**
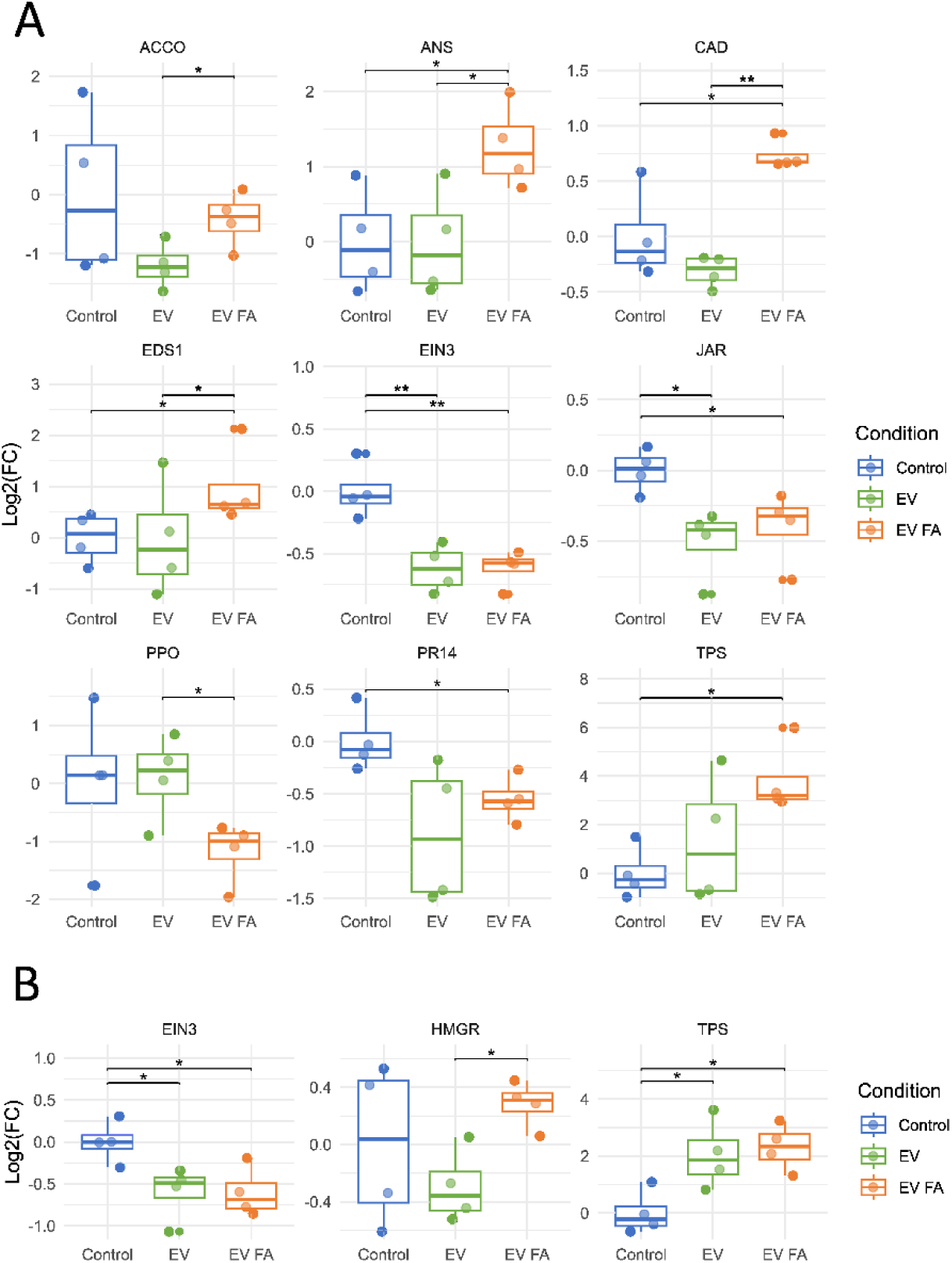
*Azospirillum* sp. B510 EVs induce systemic changes in *S. lycopersicum* defense genes expression. Transcript accumulation was determined on leaves tissues by qRT-PCR at 18 hpi **(A)** and 5 dpi **(B)** of the EV, EV FA or Control condition. Gene transcript levels were normalized using three reference genes (*TuA*, *Actin* and *GAPDH*) as internal controls. Results are expressed as the Log2-fold increase in transcript level compared to the control. *ACCO*: 1 aminocyclopropene-1-carboxylate oxidase; *ANS*: anthocyanidin synthase; *CAD*: cinnamyl alcohol dehydrogenase; *EDS1*: disease resistance protein EDS1; *EIN3*: EIN3-Binding F Box Protein 1; *HMGR*: hydroxymethyl glutarate-CoA reductase; *JAR*: jasmonate resistant 1; *PPO*: polyphenol oxidase; *PR14*: pathogenesis-related protein 14; *TPS*: (E,E)-α-farnesene synthase. Results show the mean Log2(FC) of each defense gene expression (n = 4, 15 plants/pool, Tukey analysis, * *pv* > 0.05, ** *pv* > 0.01).

**Figure 6:**
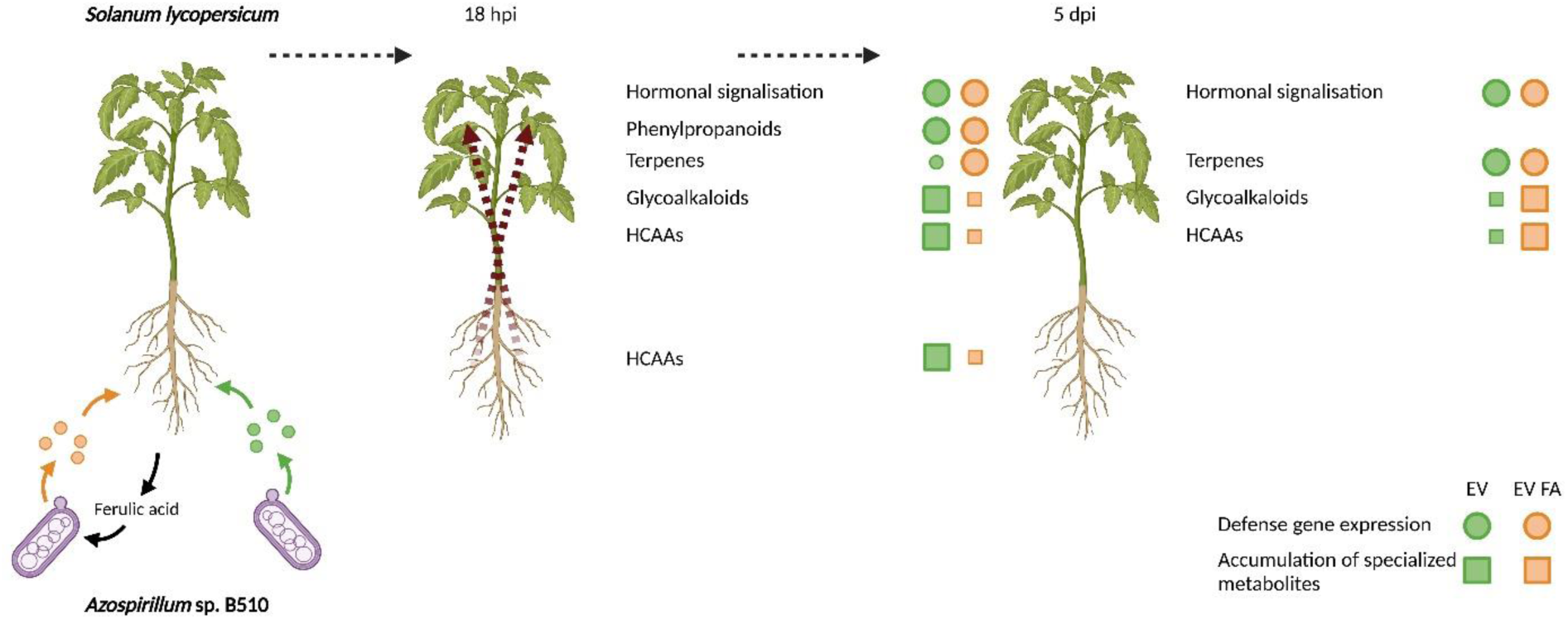
Systemic effects of *Azospirillum* sp. B510 EVs on *S. lycopersicum*. Effect of *Azospirillum* sp. B510 EVs extracted from ABS (EV - green) and ABS containing FA (EV FA - orange) medium on *S. lycopersicum* physiology. The impact of EVs on local and systemic defense gene expression (circle) and over-abundanceof specialized metabolites (square) was observed 18h and 5 days post-inoculation (18 hpi and 5 dpi).

## Discussion

Bacterial extracellular vesicles (EVs) are emerging as key mediators of the interaction between phytobacteria and their plant hosts, notably due to their impact on the plant immune system. While most studies have focused on the role of these EVs in pathosystems, their functions in beneficial plant-bacteria interactions are less explored (Zhou *et al*., 2022). In this study, we investigate how the presence of ferulic acid affects the cargo of EVs produced by the phytobeneficial bacterium *Azospirillum* sp. B510 and assess their impact on the plant partner.

### Plant environment influences bacterial EVs cargoes

*Azospirillum* sp. B510 produces EVs of a size within the range described for phytobacteria (20-400 nm) (Janda and Robatzek, 2022). These *Azospirillum* sp. B510 EVs encapsulate a large protein cargo with an amount of detected proteins in accordance with a recent proteomic study on plant-associated bacteria EVs (McMillan and Kuehn, 2022). As proteins are not the sole content of bacterial EVs cargos, we also described the lipidic and metabolic composition of *Azospirillum* sp. B510 EVs to extend the current knowledge on the global composition of plant-associated bacteria EVs. Contrary to other bacterial models, and to our expectations, the addition of a plant compound in the growth medium of *Azospirillum* sp. B510 only impacted the proteinaceous cargoes of its EVs (Li *et al*., 2022).

The presence of sub-lethal doses of ferulic acid in the environment of several phytobacteria induces the production of enzymes that allow the degradation of FA into protocatechuic acid (PCA) through either the CoA-dependent ꞵ-oxidation or the CoA-dependent non-ꞵ-oxidation pathway (Parke and Ornston, 2003; Hirakawa *et al*., 2012; Campillo *et al*., 2014; Lowe *et al*., 2015; Calero *et al*., 2018; Meyer *et al*., 2018; Chen *et al*., 2022). The PCA is then degraded by enzymes from the *pca* operon and converted through the ꞵ-ketoadipate pathway into succinyl-CoA and acetyl-CoA that are integrated in the tricarboxylic acid cycle (Parke, 1995; Wang *et al*., 2015). Here, we show that *Azospirillum* sp. B510 is able to degrade FA and to load its EVs with enzymes belonging to the CoA-dependent ꞵ-oxidation pathway (AZL_b03680 and AZL_b03690) and to the PCA degradation pathway (AZL_d00460, AZL_d00470, AZL_d00480). These enzymes correspond to the end of the CoA-dependent ꞵ-oxidation pathway and the beginning of the PCA degradation pathway. This is consistent with the timing of EVs extraction, as at this point, FA or VA was no longer detected in the supernatant. These results are in agreement with those obtained for *P. putida* KT2440 where EVs were shown to be enriched in enzymes of the CoA-dependent ꞵ-oxidation pathway and PCA degradation pathway when grown in lignin-rich medium (Salvachúa *et al*., 2020). In addition to the PCA degradation activity shown in *P. putida* EVs, our results provide a further example of the influence of the plant on the proteinaceous cargoes of phytobacteria EVs and point toward the idea that EVs could serve as remote factories of aromatic monomers degradation, thus playing a strategic role in nutrient acquisition in the plant environment.

Among the other proteins modulated by FA in *Azospirillum* sp. B510 EVs, flagellar proteins were found to be depleted in the FA condition. As in our study, the presence of flagellar proteins in EVs cargoes has already been described even when no flagellar apparatus was observed in electron microscopy (Jonca *et al*., 2021). Thus, these proteins were most likely components of the EVs cargoes and not the product of co-purifications between EVs and flagellar debris (Bahar *et al*., 2016; Janda *et al*., 2023). While EF-Tu, a well-described MAMP, was found in EVs cargo from both conditions, FliC was the only known proteinaceous MAMP to be modulated by the presence of FA. The application of *Azospirillum* sp. B510 EVs (both EV and EV FA) induced a strong systemic response in *S. lycopersicum*. However, as demonstrated previously on *Arabidopsis thaliana*, plant response to phytobacterial EVs is complex and cannot be attributed to the sole presence of FliC and/or EF-Tu (McMillan *et al*., 2021; Chalupowicz *et al*., 2023; Janda *et al*., 2023). Indeed, the application of *Xanthomonas campestris* pv. *campestris* EVs on *A. thaliana* was shown to induce a MAMP-independent response but a potentially EVs lipid composition-dependent response (Tran *et al*., 2022). The environment in which bacteria grow was shown to modify the EVs lipidic composition in *Sinorhizobium fredii* (Li *et al*., 2022). We thought that FA might also induce changes in the composition of the bacterial membrane and consequently on the lipidic composition of *Azospirillum* sp. B50 EVs. Despite no observed effect of FA on the lipidic composition of EVs, we observed a difference in systemic plant responses between EV and EV FA, suggesting the implication of other compounds of EVs cargoes. In any case, the plant reacted to *Azospirillum* sp. B510 EVs. According to Tran et *al.*, 2022, the difference in fatty acids and phospholipid composition between the bacterial EVs and the plant cell membrane combined with the physical insertion of EVs in the membrane is sufficient to trigger a plant immune response (Tran *et al*., 2022).

### Bacterial EVs modulate the physiology of the plant

*Azospirillum* sp. B510 is a phytobeneficial bacteria that can colonize the roots of tomato and rice and induce a systemic resistance (ISR) against bacterial and fungi leaf pathogens. Surprisingly, it has been demonstrated that this ISR is not mediated by salicylic accumulation (SA) nor by Pathogenis-Related (PR) gene expression and to date the defense hormonal pathways modulated by *Azospirillum* sp. B510 remain unclear (Yasuda *et al*., 2009; Fujita *et al*., 2017). As demonstrated in this study, EVs from *Azospirillum* sp. B510 induced a systemic repression of genes from the ethylene (Et) pathway that lasted for up to 5 days. Whereas repressing genes of the jasmonate and Et pathway, *Azospirillum* sp. B510 EV FA induced the expression of genes of the SA pathway, as described in *A. thaliana* following the contact with *P. syringae* EVs (McMillan *et al*., 2021). While the authors suggest that plants sense the EVs differently if they originate from pathogens or beneficial bacteria, our results show that the distinction between beneficial and pathogenic bacterial EVs by the plant could be more nebulous than previously thought. As plant defense responses are complex and cannot be assayed only by testing hormonal responses, our study aimed to broaden the defense pathways impacted by EVs. *Azospirillum* sp. B510 EVs also modulated the expression of genes involved in the biosynthesis of HCAs and steroidal alkaloids, two classes of molecules involved in plant defense against pathogens (Mandal *et al*., 2010; Panda *et al*., 2022).

To date, published studies mainly focused on the effect of phytobacterial EVs on plants by assessing their impact on the local expression of defense genes, on the accumulation of targeted defense hormonal compounds and on the ROS burst assay (Janda and Robatzek, 2022). While these methods are highly relevant, plants produce a wide range of primary and secondary metabolites in response to a bacterial presence, making metabolomic studies a pertinent way to decipher plant-microorganisms interactions (Adeniji *et al*., 2020). Indeed, they complement other omics methods by identifying key molecular markers involved in plant response to microorganisms. Here, we show for the first time that phytobacterial EVs induce both local and systemic responses on the plant metabolome. More precisely, EVs from *Azospirillum* sp. B510 impacted metabolites belonging to the Hydroxycinnamic Acids Amides derivatives (HCAAs) in roots and to steroidal alkaloids in aerial parts, differently depending on their cargo content, in accordance with the pattern of plant gene expression.

HCAs and HCAAs were shown to accumulate in presence of pathogens as they possess antimicrobial activities and are used to strengthen the plant cell wall (Borges *et al*., 2013; Zeiss *et al*., 2021). *Azospirillum* sp. B510 was previously shown to induce the over-abundance of two HCAAs (i.e. *N*-*p*-coumaroylputrescine and *N*-feruloyltyramine) in rice roots 7 days after inoculation (Valette *et al*., 2020). We showed that *Azospirillum* sp. B510 EVs have an effect on 8 HCAAs of *S. lycopersicum* roots but this effect was resorbed 5 days after inoculation. More precisely, EVs coming from a plant-mimicking environment (EV FA) were shown to significantly lower the abundance of two FA derivatives compared to EVs extracted from a control condition (EV) and this trend was observed for the other 6 annotated HCAAs. In the Solanaceae family, glycoalkaloids are defense molecules mainly found in stems, leaves and fruits that act as pests deterrent and a chemical barrier against microbial pathogens (Keukens *et al*., 1995; Friedman, 2002; Milner *et al*., 2011). ɑ-tomatine, one of the most studied glycoalkaloid in tomato, has been shown to be over-abundant in leaves of *S. lycopersicum* after inoculation of the phytopathogen *Ralstonia solanacearum* (Zeiss *et al*., 2019). *Azospirillum* sp. B510 EVs on its own induced the over-abundance of ɑ-tomatine isomers, hydroxytomatine isomers and dehydrotomatine isomers but this was not observed with EV FA.

EVs from the plant-mimicking condition seem to lower the plant defenses to which they have been applied. Other studies reported an increase in plant defense potentiation by microbial EVs (Bahar *et al*., 2016; Chalupowicz *et al*., 2023; Janda *et al*., 2023). However, contrary to our study, they used EVs extracted from laboratory growth media. Our results strengthen the idea of using growth conditions specifically dedicated to the study of plant-microbes interactions.

## Conclusion: EVs-mediated interactions are dynamic

In this study, we show that EVs from the phytobenefical bacterium *Azospirillum* sp. B510 can deeply impact plant physiology, especially its defense mechanism through the modulation of the expression of defense genes and therefore the over-abundance of plant specialized metabolites involved in the protection against pathogens. To our knowledge, this is the first evidence of a plant systemic response, as in roots to leaves, in response to bacterial EVs. This study brings new insights on an existing dynamism mediated by bacterial EVs in plant-bacteria interactions. Given the importance of the environment on the composition of bacterial EVs and their effects on the plant partner, it seems crucial to use host-mimicking media in order to understand the full complexity of the role of EVs in host-bacteria interactions.

## Author Contribution

TZP, LV and CL designed the research; TZP, LV and CL wrote the manuscript; FWD and GC proof-read the manuscript. TZP, LD, VG and IK performed the *S. lycopersicum* metabolites extraction and analysis. PYD performed the Cryo-electron microscopy observations. MG performed the *S. lycopersicum* defense gene expression analysis. FB helped to design the EVs extraction protocole. TZP carried out all the other experiments and statistical analysis and prepared the figures.

## Acknowledgments

We acknowledge the DTAMB platform of BioEEnViS Research Federation for providing access to the ultracentrifuge; the CTµ (Centre Technologique des Microstructures, University Claude Bernard Lyon 1) for the electronic microscopy observations; the Plateforme Genomique Environnementale (UMR 5557) for their help on protein experimentation; the Centre d’Etudes des Substances Naturelles (UMR 5557) for their help and recommendations on metabolomic experiment, analysis and fruitful discussions (M. Rey, G. Meiffren, P-E Mercier and P. Vergne)

We acknowledge A. Page from Protein Science Facility of SFR Biosciences (UAR3444/CNRS, US8/Inserm, ENS de Lyon, UCBL) for mass spectrometry analysis. We acknowledge F-X. Gillet and I. Tguafaiti (MAP laboratory, UMR 5240 University Claude Bernard Lyon 1-INSA-Bayer) for kindly providing access to the NTA. We would like to thank A. Lautrey for his original ideas for the plant growing system and B. Schaack for providing liposomes stocks. We acknowledge F. Nazaret for her help on proof-reading the manuscript.

Graphical figures were created using Biorender.com

