## Supplementary_Figures_and_Tables for "Extracellular vesicles of a phytobeneficial bacterium trigger distinct systemic response in plant"

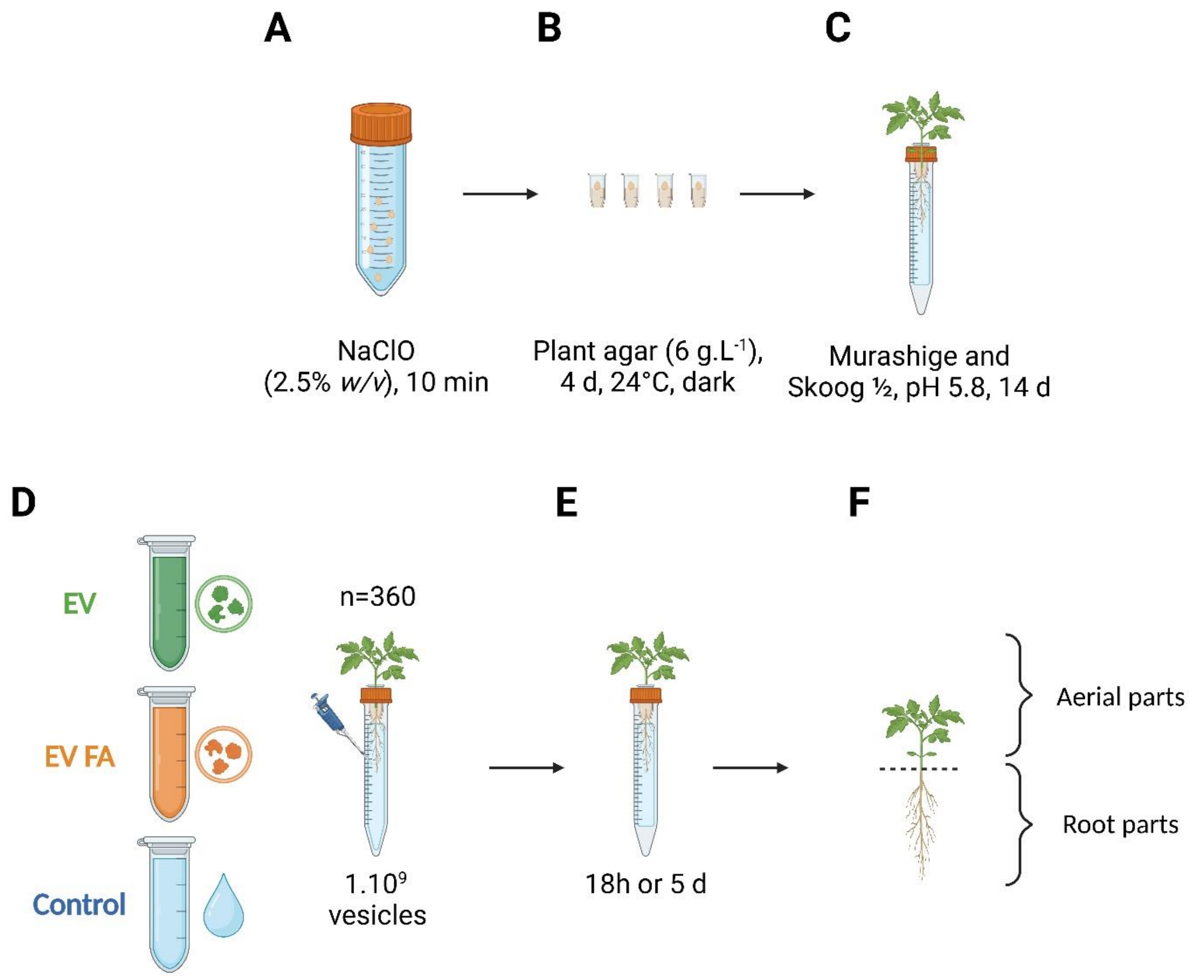


**Supplementary Figure 1:** Experimental workflow design of growth conditions of *S. lycopersicum* and root inoculation**. (A)** Seed disinfection, **(B)** germination and **(C)** seedling growth. **(D)** Application of *Azospirillum* sp. B510 EV, EV FA or water (Control condition) on roots of tomato seedling **(E)** and growth for 18h or 5 days before **(F)** plant tissues harvesting. Plant tissues were therefore used for analyses of plant metabolites accumulation and defense gene expression.

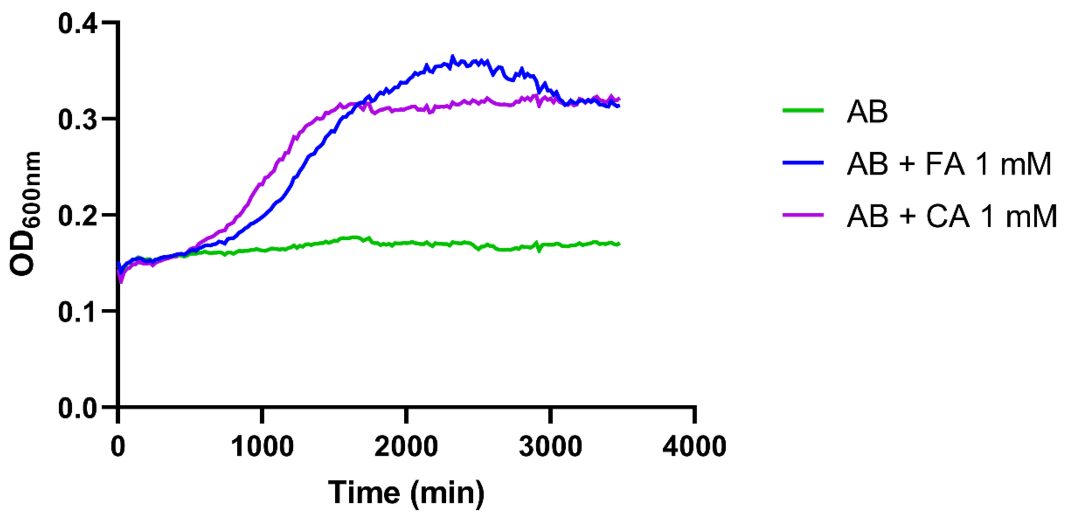


**Supplementary Figure 2:** Growth analysis of *Azospirillum* sp. B510 with HCA as the sole carbon source. *Azospirillum* sp. B510 was grown in microplates in AB minimal medium supplemented with either ferulic acid (1 mM) or coumaric acid (1 mM). FA: Ferulic acid, CA: Coumaric acid. Analyses were conducted on 3 biological replicates.

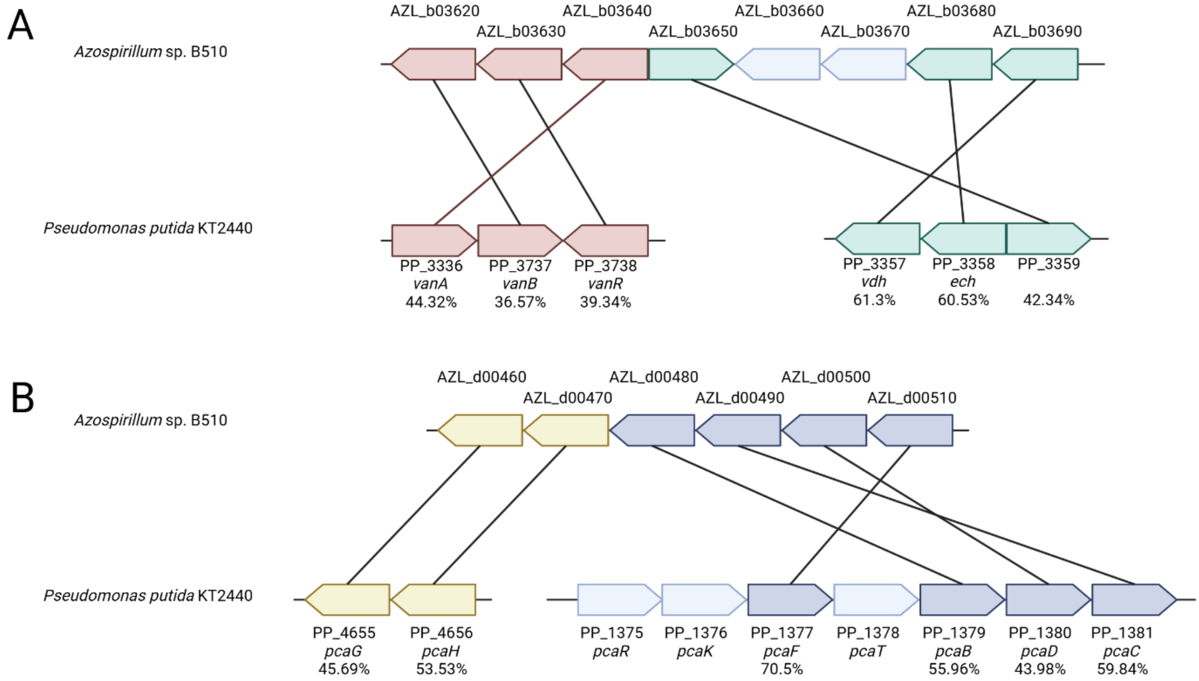


**Supplementary Figure 3:** Synteny of **(A)** HCA degradation operon and **(B)** PCA degradation operon between *Azospirillum* sp. B510 and *Pseudomonas putida* KT2440. Values represent the percentage of identity between *Azospirillum* sp. B510 and *Pseudomonas putida* KT2440 genes.

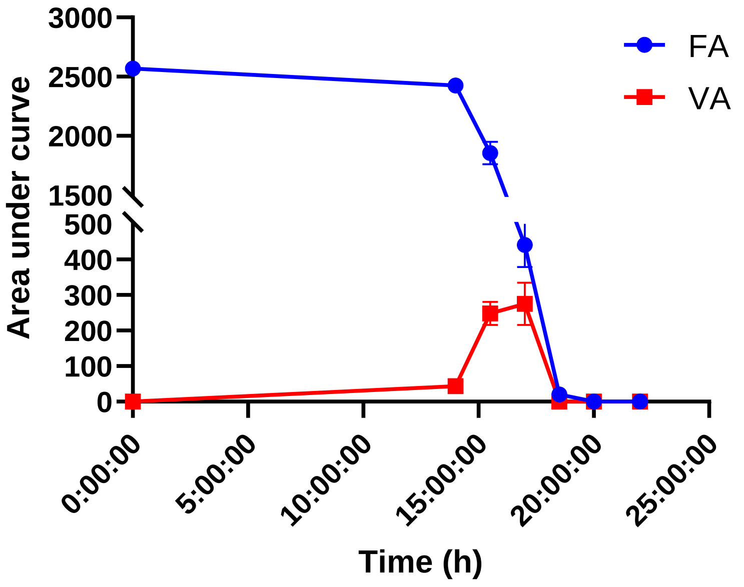


**Supplementary Figure 4:** Degradation of ferulic acid by *Azospirillum* sp. B510 and apparition of vanillic acid in the growth medium (AB minimal medium). FA: Ferulic acid. VA: Vanillic acid. Analyses were conducted on 3 biological replicates.

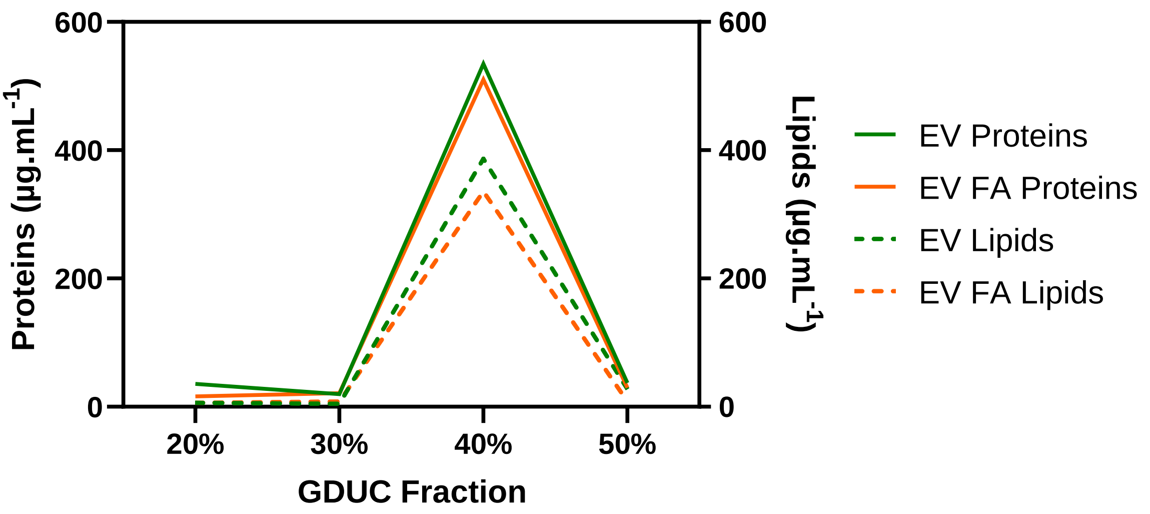


**Supplementary Figure 5:** Proteins and lipids quantification in the gradients of Optiprep ultracentrifuge used to purify EVs for subsequent experiments. Proteins were dosed using the BCA assay and lipids were dosed using the FM4-64 assay. These data are representative of every EVs purification done in the present study.

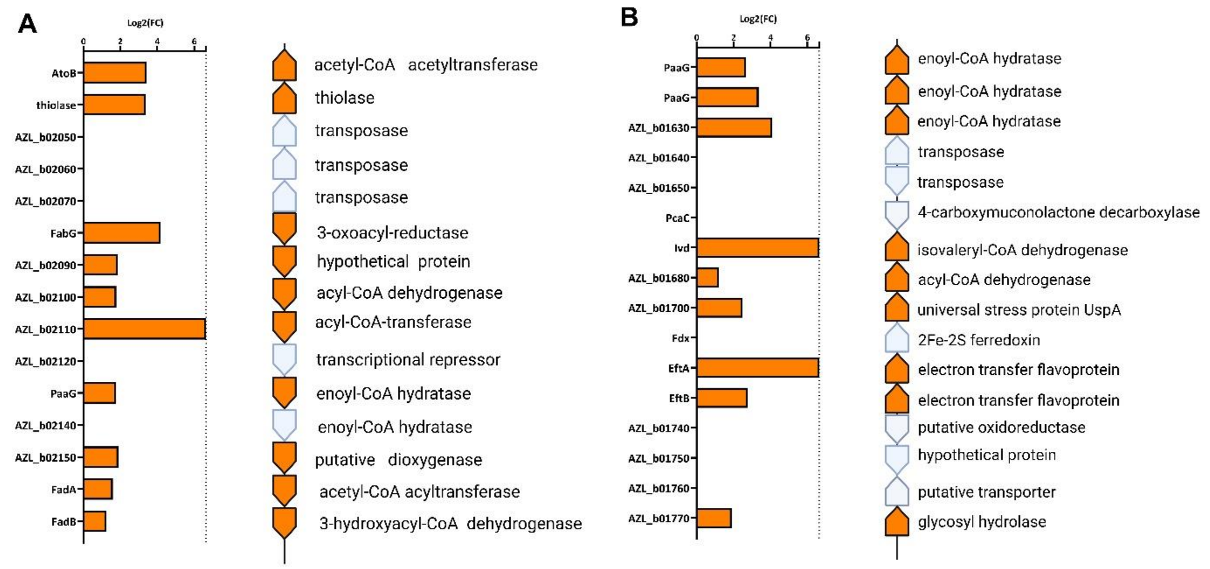


**Supplementary Figure 6:** Genes clusters coding for proteins enriched in *Azospirillum* sp. B510 EVs by the presence of FA in the environment. Genes clusters located on the replicon B of *Azospirillum* sp. B510 coding for **(A)** 10 proteins and **(B)** 9 proteins. Proteins in orange are enriched in *Azospirillum* sp. B510 EVs in presence of FA.

A

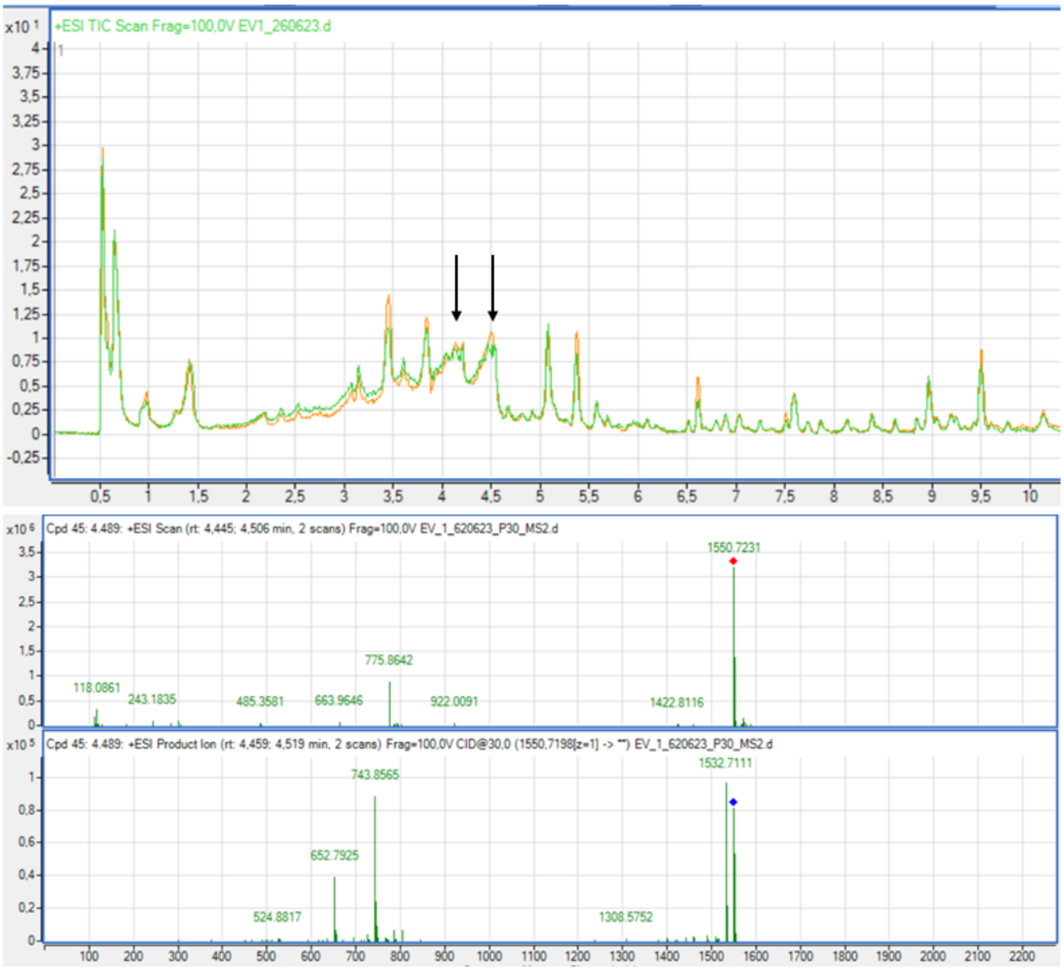


B

**Supplementary Figure 7:** Iodixanol (used for ultracentrifuge gradient preparation) ions found in *Azospirillum* sp. B510 EV samples. **(A)** Total Ion Current chromatogram (ESI+) obtained by UHPLC-ESI-MS QTOF analysis of EV sample extract. Peaks with arrows indicate iodixanol ions. **(B)** MS spectrum and MS/MS spectrum (CID+30 eV) of iodixanol.

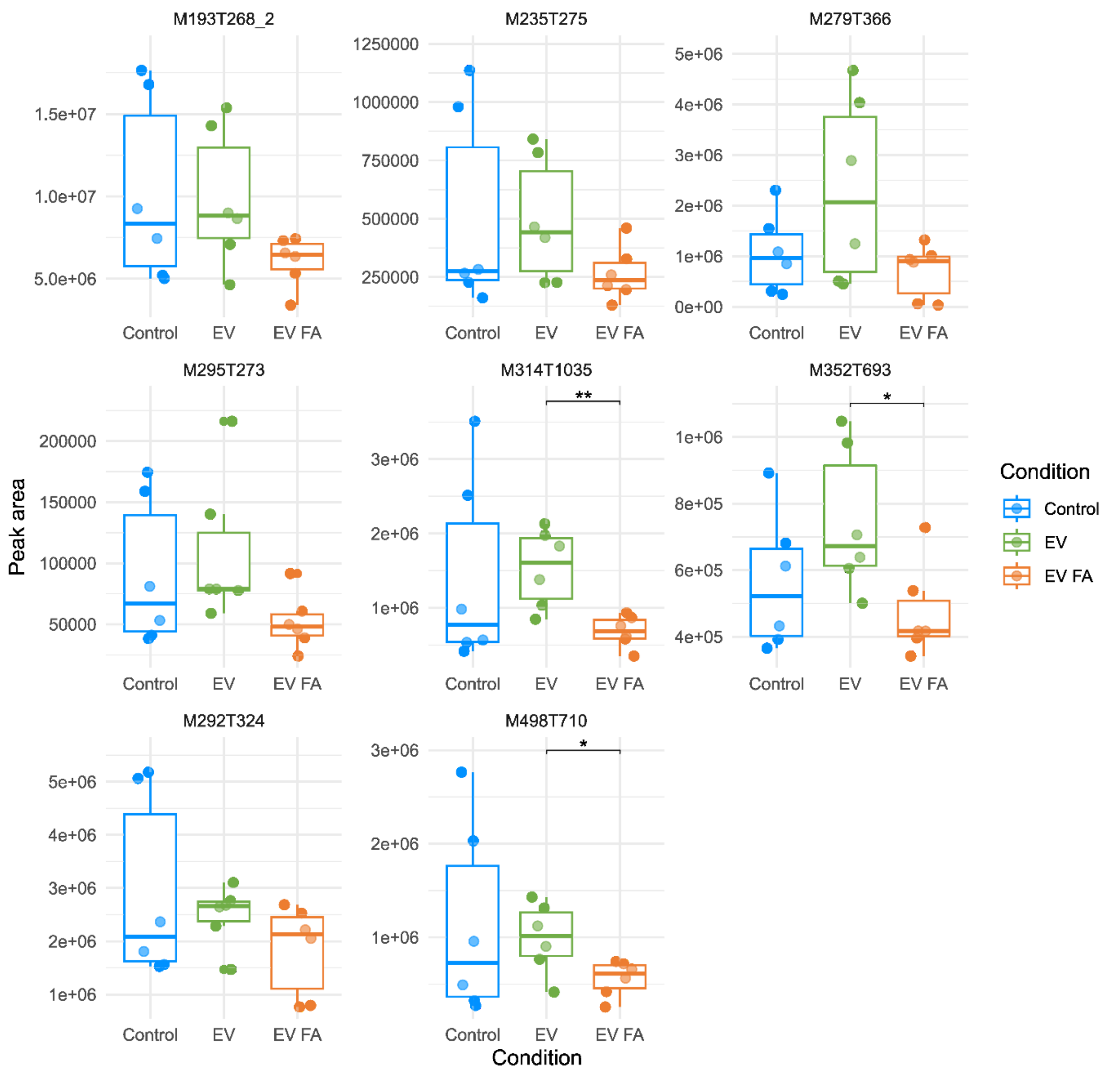


**Supplementary Figure 8:** Impact of *Azospirillum* sp. B510 EVs on the HCAAs content of *S. lycopersicum* roots 18 hpi. M193T268_2 = *N*-benzoyl-putrescine; M295T273 = *N*-sinapoylputrescine; M235T275 = *N*-(*p*-coumaroyl)-putrescine; M292T324 = *N*-(*p*-coumaroyl)-spermidine; = M279T366 = *N*-feruloylcadaverine; M352T693 = *N*-cis-feruloyloctopamine; M498T710 = *N*-(feruloyl-O-hexoside)-tyramine and M314T1035 = *N*-trans-isoferuloyltyramine. Results show the mean concentration of each ion (n = 6, 10 plants/pool, Tukey analysis, * *pv* > 0.05, ** *pv* > 0.01).

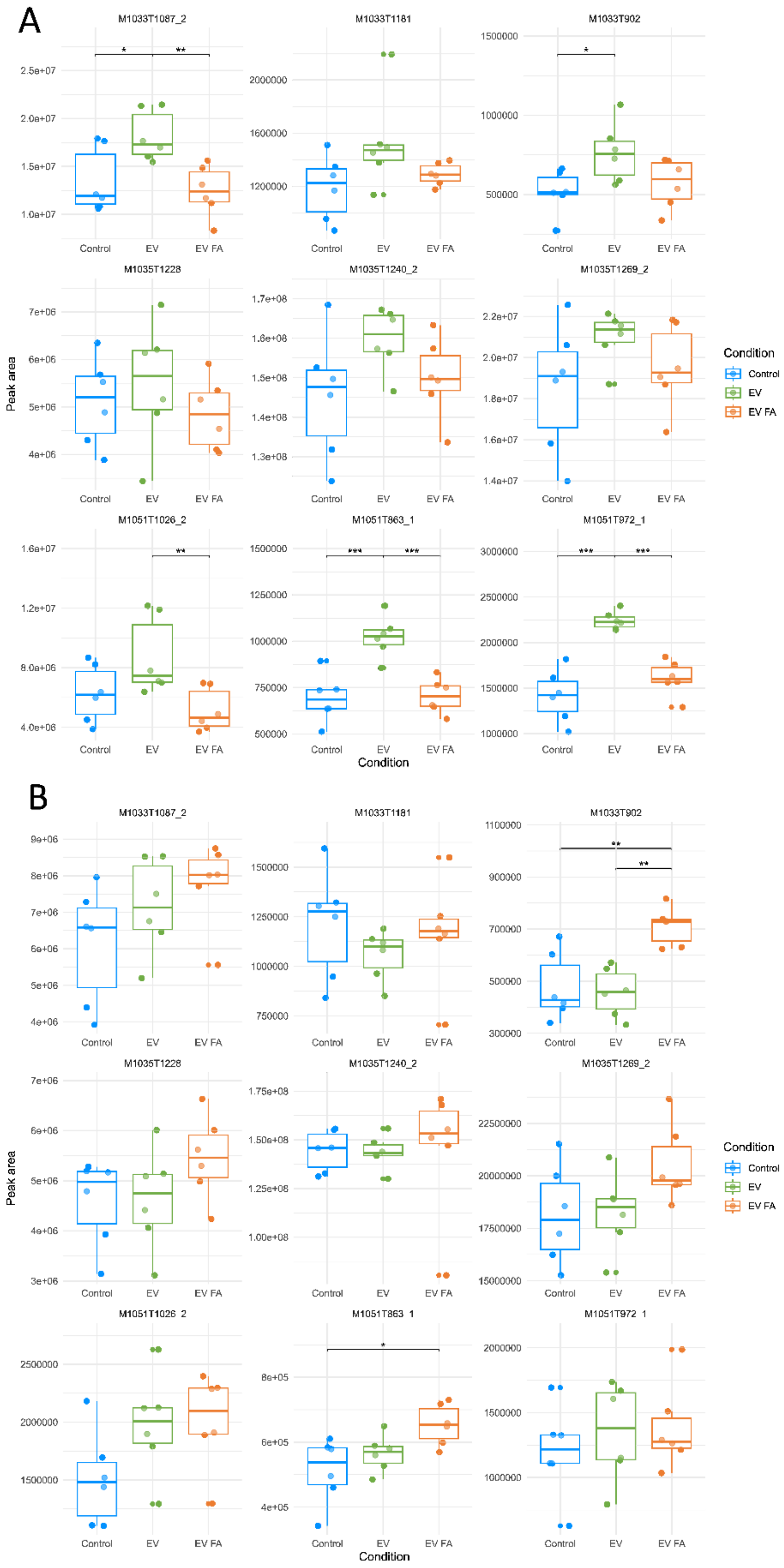


**Supplementary Figure 9:** Impact of *Azospirillum* sp. B510 EVs on the glycoalkaloid content of *S. lycopersicum* aerial parts18 hpi **(A)** and 5 dpi **(B)**. M1033T902, M1033T1087_2, M1033T1181 = dehydrotomatine isomers; M1035T1240_2 = tomatine; M1035T1228, M1035T1269 = tomatine isomers; M1051T863_1, M1051T972_1, M1051T1026_2 = hydroxytomatine isomers. Results show the mean concentration of each ion (n = 6, 10 plants/pool, Tukey analysis, * *pv* > 0.05, ** *pv* > 0.01, *** *pv* > 0.001).

**Supplementary Table 1**: Top 200 most abundant proteins in *Azospirillum* sp. B510 extracellular vesicles ranked by abundance. Table was depleted from 52 ribosomal proteins. (a) Protein cellular localization prediction by CELLO (http://cello.life.nctu.edu.tw/).

| Accession | Gene ID | Description | Gene name | # Unique Peptides | MW [kDa] | Predicted location (a) |
|---|---|---|---|---|---|---|
| D3P0K4 | AZL_a06520 | Flagellin | | 42 | 64,8 | Extracellular |
| D3NRF5 | AZL_003460 | Chaperonin GroEL | *hspD1* | 58 | 57,6 | Cytoplasmic |
| D3NSN9 | AZL_007800 | Iron complex outermembrane recepter protein | | 61 | 74,4 | OuterMembrane |
| D3NWM9 | AZL_021700 | Ketol-acid reductoisomerase | *ilvC* | 35 | 37,3 | Cytoplasmic |
| D3NSH6 | AZL_007170 | Flagellin | | 35 | 64,5 | Extracellular |
| D3NS24 | AZL_005650 | Elongation factor Tu | *tufA* | 32 | 43 | Cytoplasmic |
| D3NVY1 | AZL_019220 | Glutamine synthetase | *glnA* | 37 | 51,8 | Cytoplasmic |
| D3NVT9 | AZL_018800 | Elongation factor G | *fusA* | 75 | 78,9 | Cytoplasmic |
| D3P8D6 | AZL_f01550 | Calcium-binding protein | | 8 | 268,8 | Extracellular |
| D3NT74 | AZL_009650 | Fructose-bisphosphate aldolase class I | *fbaB* | 27 | 33,2 | Cytoplasmic |
| D3NUM6 | AZL_014670 | Uncharacterized protein | | 16 | 41,4 | Extracellular |
| D3NRA6 | AZL_002970 | Pyruvate, phosphate dikinase | *ppdK* | 65 | 97,8 | Cytoplasmic |
| D3NQK0 | AZL_000410 | Chaperone protein DnaK | *dnaK* | 57 | 68,3 | Cytoplasmic |
| D3NS31 | AZL_005720 | DNA-directed RNA polymerase subunit beta | *rpoB* | 117 | 155 | Cytoplasmic |
| D3P0B8 | AZL_a05660 | Malate dehydrogenase | *maeB* | 55 | 82,2 | Cytoplasmic |
| D3NXT4 | AZL_025750 | Malate dehydrogenase | *mdh* | 22 | 33,9 | Cytoplasmic |
| D3NXJ4 | AZL_024850 | Uncharacterized protein | | 32 | 29,9 | Cytoplasmic |
| D3NUS6 | AZL_015170 | Dihydrolipoyl dehydrogenase | *pdhD* | 44 | 49,1 | Cytoplasmic |
| D3NS32 | AZL_005730 | DNA-directed RNA polymerase subunit beta' | *rpoC* | 113 | 154,1 | Cytoplasmic |
| D3P1D1 | AZL_a09290 | Transcriptional regulator | | 4 | 25,4 | Cytoplasmic |
| D3NX05 | AZL_022960 | Uncharacterized protein | | 21 | 36,8 | Extracellular |

| | | | | | | |
|---|---|---|---|---|---|---|
| D3P897 | AZL_f01160 | NAD-dependent epimerase/dehydratase | | 11 | 35 | Cytoplasmic |
| D3NYT7 | AZL_a00350 | Outer membrane protein | | 14 | 20,7 | Periplasmic |
| D3NX22 | AZL_023130 | Acetyl-CoA C-acetyltransferase | *atoB* | 28 | 40,3 | Cytoplasmic |
| D3NXT5 | AZL_025760 | Succinate--CoA ligase subunit beta | *sucC* | 25 | 42,4 | Cytoplasmic |
| D3NRU0 | AZL_004810 | Iron complex outermembrane recepter protein | | 59 | 78,2 | OuterMembrane |
| D3NV18 | AZL_016090 | Citrate synthase | *gltA* | 30 | 49 | Cytoplasmic |
| D3NXY5 | AZL_026260 | Porin domain-containing protein | | 27 | 41 | OuterMembrane |
| D3NXY6 | AZL_026270 | Porin domain-containing protein | | 25 | 40 | OuterMembrane |
| D3NQQ8 | AZL_000990 | Glutamate synthase large chain | *gltB* | 92 | 165,6 | Cytoplasmic |
| D3NXT8 | AZL_025790 | Dihydrolipoyllysine-residue succinyltransferase | *sucB* | 25 | 42,8 | Cytoplasmic |
| D3P5Y7 | AZL_d02400 | Beta_helix domain-containing protein | | 55 | 110,9 | Extracellular |
| D3NV92 | AZL_016830 | Isocitrate dehydrogenase | *icd* | 37 | 45,7 | Cytoplasmic |
| D3NZB2 | AZL_a02100 | PilS domain-containing protein | | 4 | 21,9 | Extracellular |
| D3NYM2 | AZL_028630 | Polyribonucleotide nucleotidyltransferase | *pnp* | 47 | 76,8 | Cytoplasmic |
| D3NWF2 | AZL_020930 | Inosine-5'-monophosphate dehydrogenase | *guaB* | 43 | 52,2 | Cytoplasmic |
| D3NQK8 | AZL_000490 | Transcriptional regulatory protein | | 11 | 15,1 | Periplasmic |
| D3NR20 | AZL_002110 | Argininosuccinate synthase | *argG* | 39 | 45,8 | Cytoplasmic |
| D3NUS9 | AZL_015200 | Pyruvate dehydrogenase E1 component subunit beta | *pdhB* | 27 | 49,4 | Cytoplasmic |
| D3NUT0 | AZL_015210 | Pyruvate dehydrogenase E1 component subunit alpha | *pdhA* | 27 | 37,1 | Cytoplasmic |
| D3P6R0 | AZL_d05130 | Beta_helix domain-containing protein | | 33 | 67,9 | Extracellular |
| D3NXK3 | AZL_024940 | ATP synthase subunit alpha | *atpA* | 39 | 54,7 | Cytoplasmic |
| D3NRY8 | AZL_005290 | Phosphomethylpyrimidine synthase | *thiC* | 44 | 62,9 | Cytoplasmic |
| D3NV34 | AZL_016250 | Elongation factor Ts | *tsf* | 24 | 32,3 | Cytoplasmic |

| | | | | | | |
|---|---|---|---|---|---|---|
| D3NXT7 | AZL_025780 | 2-oxoglutarate dehydrogenase E1 component | *sucA* | 57 | 109,3 | Cytoplasmic |
| D3NVD4 | AZL_017250 | Phasin | | 17 | 15,4 | Cytoplasmic |
| D3NUK2 | AZL_014430 | Homospermidine synthase | | 28 | 52,3 | Cytoplasmic |
| D3P3W6 | AZL_c00520 | Outer membrane protein | | 19 | 26,2 | OuterMembrane |
| D3NW08 | AZL_019490 | Probable cytosol aminopeptidase | *pepA* | 36 | 52,6 | Cytoplasmic |
| D3NRJ0 | AZL_003810 | Chaperone protein HtpG | *hsp90A* | 52 | 69,4 | Cytoplasmic |
| D3P8B0 | AZL_f01290 | Cadherin domain-containing protein | | 61 | 405,9 | Extracellular |
| D3NY11 | AZL_026520 | Adenosylhomocysteinase | *ahcY* | 29 | 47,2 | Cytoplasmic |
| D3NQH9 | AZL_000200 | Malate dehydrogenase | *maeB* | 52 | 81,7 | Cytoplasmic |
| D3NXK7 | AZL_024980 | Probable transaldolase | *talB* | 19 | 23,3 | Cytoplasmic |
| D3P5I6 | AZL_d00890 | Alkyl hydroperoxide reductase C | *ahpC* | 15 | 20,6 | Cytoplasmic |
| D3NWT9 | AZL_022300 | 2-isopropylmalate synthase | *leuA* | 51 | 61,4 | Cytoplasmic |
| D3NR71 | AZL_002620 | Carbamoyl-phosphate synthase large chain | *carB* | 67 | 115,5 | Cytoplasmic |
| D3P0P1 | AZL_a06890 | Endoglucanase | | 40 | 87,6 | Extracellular |
| D3NRU5 | AZL_004860 | Tyrosine--tRNA ligase | *tyrS* | 27 | 45,4 | Cytoplasmic |
| D3P359 | AZL_b05660 | TRAP dicarboxylate transporter | | 29 | 35,4 | Periplasmic |
| D3NTG7 | AZL_010580 | RNA polymerase sigma factor RpoD | *rpoD* | 34 | 75,3 | Cytoplasmic |
| D3P0A3 | AZL_a05510 | Flagellar basal body protein | *flgE* | 29 | 89,6 | Extracellular |
| D3NQY4 | AZL_001750 | Succinate dehydrogenase flavoprotein subunit | *sdhA* | 30 | 64,9 | Cytoplasmic |
| D3P505 | AZL_c04410 | Aconitate hydratase B | *acnB* | 64 | 92,5 | Cytoplasmic |
| D3NSZ9 | AZL_008900 | 3-isopropylmalate dehydrogenase | *leuB* | 30 | 40,2 | Cytoplasmic |
| D3NUS8 | AZL_015190 | Acetyltransferase component of pyruvate dehydrogenase complex | *pdhC* | 24 | 45,2 | Periplasmic |
| D3P008 | AZL_a04560 | Alanine--tRNA ligase | *alaS* | 68 | 95,7 | Cytoplasmic |

| | | | | | | |
|---|---|---|---|---|---|---|
| D3P0A1 | AZL_a05490 | Uncharacterized protein | | 46 | 72,6 | OuterMembrane |
| D3NVD3 | AZL_017240 | Flagellin domain protein | | 28 | 64,3 | Extracellular |
| D3NVX3 | AZL_019140 | Lon protease | *lon* | 54 | 89,2 | Cytoplasmic |
| D3NXL1 | AZL_025020 | Dihydrolipoyl dehydrogenase | *pdhD* | 30 | 48,7 | Cytoplasmic |
| D3NWN3 | AZL_021740 | Acetolactate synthase | *ilvB* | 32 | 63,9 | Cytoplasmic |
| D3NUC0 | AZL_013610 | Periplasmic serine endoprotease DegP-like | | 26 | 58,5 | Periplasmic |
| D3NRR6 | AZL_004570 | DUF4139 domain-containing protein | | 40 | 79,8 | OuterMembrane |
| D3NY30 | AZL_026710 | Enoyl-CoA hydratase | *paaG* | 18 | 27,5 | Cytoplasmic |
| D3NTK3 | AZL_010940 | Fructose-1,6-bisphosphatase | *glpX* | 23 | 34,8 | Cytoplasmic |
| D3P166 | AZL_a08640 | Autotransporter domain-containing protein | | 10 | 49,6 | OuterMembrane |
| D3NXT6 | AZL_025770 | Succinate--CoA ligase subunit alpha | *sucD* | 15 | 29,8 | Cytoplasmic |
| D3P887 | AZL_f01060 | GDP-mannose 4,6-dehydratase | *gmd* | 31 | 40,8 | Cytoplasmic |
| D3P5B7 | AZL_d00200 | Dihydroxy-acid dehydratase | *ilvD* | 45 | 65,9 | Cytoplasmic |
| D3NT81 | AZL_009720 | Transketolase | *tktB* | 35 | 70,7 | Cytoplasmic |
| D3NYL2 | AZL_028530 | S-adenosylmethionine synthase | *metK* | 31 | 41,4 | Cytoplasmic |
| D3NXK1 | AZL_024920 | ATP synthase subunit beta | *atpD* | 28 | 50 | Cytoplasmic |
| D3NQS3 | AZL_001140 | Aspartokinase | *lysC* | 33 | 43,7 | Cytoplasmic |
| D3NZY1 | AZL_a04290 | ATP-dependent Clp protease proteolytic subunit | *clpP* | 17 | 23,2 | Cytoplasmic |
| D3NTB8 | AZL_010090 | Serine hydroxymethyltransferase | *glyA* | 24 | 48,2 | Cytoplasmic |
| D3NYL8 | AZL_028590 | Translation initiation factor IF-2 | *infB* | 53 | 102,9 | Cytoplasmic |
| D3NT80 | AZL_009710 | Glyceraldehyde-3-phosphate dehydrogenase | *gapA* | 7 | 36,1 | Cytoplasmic |
| D3P6Q1 | AZL_d05040 | Endoglucanase | | 33 | 82,6 | Extracellular |
| D3NRA4 | AZL_002950 | Vitamin B12-dependent ribonucleotide reductase | *nrdE* | 70 | 135,7 | Cytoplasmic |

| | | | | | | |
|---|---|---|---|---|---|---|
| D3P0W5 | AZL_a07630 | Fumarate hydratase class I | *fumB* | 42 | 64,4 | Cytoplasmic |
| D3P5C9 | AZL_d00320 | UDP-glucose 6-dehydrogenase | *ugd* | 29 | 47,8 | Cytoplasmic |
| D3NV36 | AZL_016270 | Aspartate--tRNA ligase | *aspS* | 48 | 68,2 | Cytoplasmic |
| D3NSC3 | AZL_006640 | Adenylosuccinate lyase | *purB* | 31 | 48,4 | Cytoplasmic |
| D3NUI0 | AZL_014210 | Alpha-1,4 glucan phosphorylase | *pyg* | 51 | 94,7 | Cytoplasmic |
| D3NV27 | AZL_016180 | Outer membrane protein assembly factor BamA | *yaeT* | 59 | 87,2 | OuterMembrane |
| D3NV81 | AZL_016720 | 3-oxoacyl-[acyl-carrier-protein] synthase 2 | *fabF* | 28 | 43,6 | Cytoplasmic |
| D3NWI4 | AZL_021250 | Proline--tRNA ligase | *proS* | 31 | 48,9 | Cytoplasmic |
| D3NX21 | AZL_023120 | Acetoacetyl-CoA reductase | *phbB* | 17 | 25,3 | Cytoplasmic |
| D3NT93 | AZL_009840 | Tol-Pal system protein TolB | *tolB* | 35 | 49,2 | OuterMembrane |
| D3P0F7 | AZL_a06050 | Rhizobiocin/RTX toxin and hemolysin-type calcium binding protein | | 18 | 172,6 | Extracellular |
| D3NTT0 | AZL_011710 | GMP synthase | *guaA* | 35 | 57 | Cytoplasmic |
| D3P4Z2 | AZL_c04280 | Fe-S cluster assembly ATP-binding protein | *sufC* | 17 | 27,4 | Cytoplasmic |
| D3P637 | AZL_d02900 | Nicotinate-nucleotide diphosphorylase | *nadC* | 19 | 29,2 | Cytoplasmic |
| D3NVT5 | AZL_018760 | DNA-directed RNA polymerase subunit alpha | *rpoA* | 26 | 37,5 | Cytoplasmic |
| D3P329 | AZL_b05360 | Uncharacterized protein | | 18 | 19,1 | Periplasmic |
| D3P5D0 | AZL_d00330 | Phosphomannomutase | *manB* | 39 | 50,1 | Cytoplasmic |
| D3NXA8 | AZL_023990 | Peptide/nickel transport system substrate-binding protein | | 29 | 59,2 | Periplasmic |
| D3P8G8 | AZL_f01870 | Mannose-1-phosphate guanylyltransferase | *manC* | 29 | 53,3 | Cytoplasmic |
| D3NWK1 | AZL_021420 | NADH-quinone oxidoreductase | *nuoG* | 44 | 74,4 | Cytoplasmic |
| D3NXV7 | AZL_025980 | Phenylalanine--tRNA ligase beta subunit | *pheT* | 43 | 85 | Cytoplasmic |
| D3NS20 | AZL_005610 | Uncharacterized protein | | 31 | 46,8 | OuterMembrane |
| D3NZG5 | AZL_a02630 | Polyhydroxyalkanoate synthase | *phaC* | 43 | 79,3 | Periplasmic |

| | | | | | | |
|---|---|---|---|---|---|---|
| D3P7Q0 | AZL_e03340 | 3-hydroxybutyryl-CoA dehydrogenase | *paaH* | 17 | 30,6 | Cytoplasmic |
| D3P6Q7 | AZL_d05100 | Pyruvate kinase | *pyk* | 24 | 50,8 | Cytoplasmic |
| D3P508 | AZL_c04440 | Formate acetyltransferase | *pflD* | 47 | 84,6 | Cytoplasmic |
| D3NRR3 | AZL_004540 | Thiazole synthase | *thiG* | 18 | 27,6 | Cytoplasmic |
| D3P7Z0 | AZL_f00090 | Calcium-binding protein | | 34 | 83,6 | Extracellular |
| D3NWZ5 | AZL_022860 | Phosphoribosylformylglycinamidine synthase subunit PurL | *purL* | 44 | 78,2 | Cytoplasmic |
| D3NQM1 | AZL_000620 | Phosphoenolpyruvate carboxykinase | *pepCK* | 38 | 67,7 | Periplasmic |
| D3NU23 | AZL_012640 | O-succinylhomoserine sulfhydrylase | *metZ* | 26 | 43,7 | Cytoplasmic |
| D3NR48 | AZL_002390 | Bifunctional purine biosynthesis protein PurH | *purO* | 39 | 56,2 | Cytoplasmic |
| D3NX84 | AZL_023750 | Cold shock protein | *cspA* | 18 | 19,8 | Extracellular |
| D3NUI1 | AZL_014220 | Aspartate-semialdehyde dehydrogenase | *asd* | 22 | 36,6 | Cytoplasmic |
| D3P4X5 | AZL_c04110 | Adenylosuccinate synthetase | *purA* | 31 | 46,4 | Cytoplasmic |
| D3NWN4 | AZL_021750 | Aminotransferase | | 23 | 43,8 | Cytoplasmic |
| D3NXQ7 | AZL_025480 | Uncharacterized protein | | 9 | 35,6 | Periplasmic |
| D3NWN8 | AZL_021790 | Enoyl-[acyl-carrier-protein] reductase | *fabI* | 22 | 27,9 | Cytoplasmic |
| D3NU65 | AZL_013060 | Outer membrane factor | | 23 | 50,3 | OuterMembrane |
| D3NXL9 | AZL_025100 | Phosphoglycerate kinase | *pgk* | 30 | 41,3 | Cytoplasmic |
| D3NRF4 | AZL_003450 | Co-chaperonin GroES | *hspE1* | 9 | 10,5 | Cytoplasmic |
| D3NSG5 | AZL_007060 | Isoleucine--tRNA ligase | *ileS* | 55 | 105,4 | Cytoplasmic |
| D3NR00 | AZL_001910 | Aminotransferase | *aspB* | 21 | 43,2 | Periplasmic |
| D3NVY4 | AZL_019250 | Glutamyl-tRNA amidotransferase subunit A | *gatA* | 31 | 52,2 | Cytoplasmic |
| D3P3Z6 | AZL_c00820 | Aldehyde dehydrogenase | | 17 | 52,7 | Cytoplasmic |
| D3NVX7 | AZL_019180 | Trigger factor | *tig* | 34 | 49,4 | Cytoplasmic |

| | | | | | | |
|---|---|---|---|---|---|---|
| D3NT22 | AZL_009130 | Energy-dependent translational throttle protein EttA | *ettA* | 45 | 61,7 | Cytoplasmic |
| D3P541 | AZL_c04770 | Propionyl-CoA carboxylase | *pccA* | 40 | 72,5 | Cytoplasmic |
| D3NZR4 | AZL_a03620 | Formyltetrahydrofolate deformylase | *purU* | 17 | 32,4 | Cytoplasmic |
| D3NZH9 | AZL_a02770 | Aspartyl/glutamyl-tRNA amidotransferase subunit B | *gatB* | 33 | 53,4 | Cytoplasmic |
| D3NRP7 | AZL_004380 | Serine protein kinase | *prkA* | 51 | 74,1 | Cytoplasmic |
| D3P830 | AZL_f00490 | NAD-dependent epimerase/dehydratase | | 27 | 34,7 | Cytoplasmic |
| D3NXI3 | AZL_024740 | Threonine--tRNA ligase | *thrS* | 43 | 72,9 | Cytoplasmic |
| D3NRE3 | AZL_003340 | SAM-dependent methyltransferase | | 20 | 43,8 | Cytoplasmic |
| D3NWN2 | AZL_021730 | Acetolactate synthase small subunit | *ilvH* | 14 | 18,1 | Cytoplasmic |
| D3NU67 | AZL_013080 | Valine--tRNA ligase | *valS* | 42 | 100,6 | Cytoplasmic |
| D3P7J3 | AZL_e02770 | DUF2059 domain-containing protein | | 15 | 32,2 | Periplasmic |
| D3NXA1 | AZL_023920 | 3-isopropylmalate dehydratase large subunit | *leuC* | 27 | 49,8 | Cytoplasmic |
| D3NQV5 | AZL_001460 | Chaperone protein ClpB | *clpB* | 62 | 95 | Cytoplasmic |
| D3NSH2 | AZL_007130 | Orotate phosphoribosyltransferase | *pyrE* | 18 | 25,1 | Cytoplasmic |
| D3NXS2 | AZL_025630 | Protein translocase subunit SecA | *secA* | 62 | 102,3 | Cytoplasmic |

**Supplementary Table 2: Comparison of fatty acid moieties of phospholipids in EVs between *Azospirillum* sp. B510 EV growth with or without FA.** Fatty acid moieties of phospholipids were analyzed by LC-MS/MS. The percentage of each fatty acid moieties is relative to the total fatty acid moieties. PE: Phosphatidylethanolamine ; PC: Phosphatidylcholine ; PG: Phosphatidylglycerol ; ND: Not detected. Values represent the mean of three biological replicates.

| | PE | | PC | | PG | |
|---|---|---|---|---|---|---|
| | **EV** | **EV FA** | **EV** | **EV FA** | **EV** | **EV FA** |
| **16:0 16:0** | 0.19 | 0.20 | ND | ND | 0.21 | 0.19 |
| **16:0 16:1** | 0.97 | 0.89 | 1.03 | 1.06 | 0.88 | 0.74 |
| **16:0 18:0** | 0.28 | 0.27 | 0.41 | 0.29 | 0.42 | 0.43 |
| **16:0 18:1** | 8.70 | 9.18 | 10.06 | 10.45 | 12.18 | 12.21 |
| **16:1 16:1** | 1.57 | 1.19 | 1.41 | 1.27 | 0.90 | 0.74 |
| **16:1 18:0** | 0.98 | 0.91 | 0.88 | 0.94 | 0.89 | 0.86 |
| **16:1 18:1** | 31.04 | 29.63 | 26.66 | 25.73 | 25.35 | 24.88 |
| **18:0 18:0** | 0.07 | 0.07 | 0.20 | 0.15 | 0.08 | 0.09 |
| **18:0 18:1** | 4.03 | 4.10 | 5.48 | 5.54 | 4.59 | 4.67 |
| **18:1 18:1** | 52.17 | 53.56 | 54.06 | 54.72 | 54.49 | 55.19 |

**Supplementary Table 3:** List of *S. lycopersicum* defense genes tested in defense gene expression analysis. Adapted from Brisset and Dugé de Bernonville, 2011.

| Defense classes and sub-classes | | Gene identifier | Gene name |
|---|---|---|---|
| Chemical or physical barriers | PR Proteins | PR-1 | Pathogenesis-related protein 1 |
| | | PR-2 | Pathogenesis-related protein 2 (glucanase) |
| | | PR-4 | Pathogenesis-related protein 4 (hevein-like) |
| | | PR-5 | Pathogenesis-related protein 5 (thaumatin-like, osmotin) |
| | | PR-8 | Pathogenesis-related protein 8 (class III chitinase) |
| | | PR-14 | Pathogenesis-related protein 14 (lipid transfer protein) |
| | | PR-15 | Pathogenesis-related protein 15 (oxalate oxidase) |
| | Phenylpropanoids pathway | PAL | Phenylalanine ammonia-lyase |
| | | CHS | Chalcone synthase |
| | | DFR | Dihydroflavonol reductase |
| | | ANS | Anthocyanidin synthase |
| | | PPO | Polyphenol oxidase |
| | Isoprenoids pathway | HMGR | Hydroxymethyl glutarate-CoA reductase |
| | | FPPS | Farnesyl pyrophosphate synthase |
| | | TPS | (E,E)-α-farnesene synthase |
| | Cysteine pathway | CSL | Cystein lyase |
| | Oxidative stress | APOX | Ascorbate peroxidase |
| | | GST | Glutathion S-transferase |
| | | POX | Peroxidase |
| | Cell-wall modifications | CALS | Callose synthase |
| | | PECT | Pectin methyl esterase |
| | | CAD | Cinnamyl alcohol dehydrogenase |
| Hormonal signalization | Salicylic acid pathway | ESD1 | Disease resistance protein EDS1 |
| | | WRKY | WRKY transcription factor 30 |
| | Jasmonic acid pathway | LOX2 | Lipoxygenase AtLOX2 |
| | | JAR | Jasmonate resistant 1 |
| | Ethylene pathway | ACCO | 1-aminocyclopropane-1-carboxylate oxidase |
| | | EIN3 | EIN3-BINDING F BOX PROTEIN 1 |
